# ”Evolution of ipsilateral breast cancer decoded by proteogenomics”

**DOI:** 10.1101/2022.07.13.499898

**Authors:** Tommaso De Marchi, Paul Theodor Pyl, Martin Sjöström, Susanne Erika Reinsbach, Sebastian DiLorenzo, Björn Nystedt, Lena Tran, Gyula Pekar, Fredrik Wärnberg, Irma Fredriksson, Per Malmström, Mårten Fernö, Lars Malmström, Johan Malmstöm, Emma Niméus

## Abstract

Ipsilateral breast tumor recurrence (IBTR) is a clinically important event, where an isolated in-breast recurrence is a potentially curable event but also associated with an increased risk of distant metastases and breast cancer death. It currently remains unclear if IBTRs are associated with molecular changes that can be explored as a resource for precision medicine strategies targeting locally recurring breast cancer. Here, we employed a recently developed proteogenomics workflow to analyze a cohort of 27 primary breast cancers and their matched IBTRs by whole genome sequencing, RNA sequencing, and mass spectrometry-based proteomics to define proteogenomic features of tumor evolution. Analysis of mutational signatures, copy number changes, and cancer specific mutations revealed a relationship with estrogen and progesterone receptor statuses and increased levels of genetic change. This in turn altered the re-programming of the transcriptome and proteome towards a recurring molecular disease phenotype with high replicating capacity and a higher degree of genomic instability possibly enhanced by high expression of *APOBEC3B*. In conclusion, this study defines how primary breast tumors differentially evolve into different ipsilateral recurrent malignancies depending on their key biomarker status, suggesting that further enhancing the genomic instability in some tumors could serve as an alternative treatment option.

## Introduction

Continuous improvements in breast cancer (BC) care has reduced the risk of local recurrences^1^. Still, about 4-11% of BCs develop a ipsilateral breast tumor recurrence (IBTR) within 10 years^2, 3^. IBTR is a clinically important event in BC, an isolated in-breast recurrence is a potentially curable event but associated with an increased risk of distant metastases (DM) and breast cancer death^4–9^. The time interval between IBTR and DM constitutes a therapeutic window to prevent further spread. Over the course of the disease, the primary tumor (PT) evolves by clonal expansion and changes in its mutational landscape. Adjuvant treatments are effective at preventing recurrent disease, but may influence the expansion of therapy resistant clones, such as *ESR1* mutations after aromatase inhibitor treatment^10, 11^. To date, limited effort has been put into characterization of IBTR molecular (e.g. DNA, RNA, protein) properties and their relation to tumor evolution and response to therapy.

The repertoire of driver and passenger mutations and their effect on the transcriptome and proteome in primary BCs has been analyzed in numerous studies. These reports have connected key drivers and tumor subtype e.g. *TP53* and *PIK3CA* mutations with estrogen receptor (ER) negative and positive tumors, respectively, as well as defined how specific mutations can impact gene expression and prognosis^12–14^, thus providing new opportunities for patient stratification and novel therapies. So far however, while investigation of DM is becoming more frequent, few studies have investigated the processes that lead to the development of IBTRs. Ultra-deep sequencing studies focusing on matched primary and distant recurrent tumors have shown the relevance of specific driver mutations such as *JAK2* in promoting tumor progression and proliferation, which has in turn catalyzed new avenues for therapeutic intervention by JAK-STAT pathway inhibition^15–17^. Genomic alterations occurring between primary and recurrent cancers, such as missense mutations and copy number (CN) changes, have further clarified mutational processes involved in the evolution to DM, such as APOBEC-mediated mutagenesis. Furthermore, recent studies have shown that both intrinsic and extrinsic factors, such as tumor subtype and treatment, specifically drive certain nucleotide changes, referred to as mutational signatures^18, 19^, which impact the progression of primary BC into DM^20^.

Here, we have employed a previously developed proteogenomics workflow^21^ to determine the evolution of IBTRs at the genomic, transcriptomic, and proteomic level from corresponding matched PTs, to investigate paths of tumor mutational evolution, identify biomarkers for therapeutic monitoring and alternative drug targets for therapy. Integrated proteogenomics analyses provide additional information regarding specific pathway activation e.g. ERBB2, and the consequent efficacy of inhibition therapy^22^, as well as an additional depth in tumor classification and biomarker selection^23–25^. Our analysis shows that the development of BC IBTRs is dependent on both hormonal receptor (ER and PgR) status of the PT, as well as changes in the DNA replication and transcription machinery in tandem with APOBEC proteins to increase genomic instability, resulting in an increased mutational load.

## Methods

### Sample cohort

Fresh frozen tumor samples (PTs and IBTRs) from 385 patients operated with BCS with and without radiotherapy in three health care regions (Southern Sweden, Uppsala-Örebro, and Stockholm) were previously collected in a multi-center cohort, previously analyzed by gene expression^26^. We selected samples based on availability for downstream DNA, RNA, and protein extraction, routine biomarkers (Estrogen Receptor, ER; Progesterone Receptor, PgR, Heregulin 2, Her2/ERBB2; proliferation marker Ki-67), follow-up information until formation of IBTR (IBTR-free survival, IBTRFS), and availability of formalin-fixed and paraffin-embedded (FFPE) material for re-analysis. A total of 54 samples (27 PTs matched by 27 IBTRs) was selected (**Table S1**). Samples for germline DNA whole genome sequencing (WGS) were available for two patients (S12 and S18). All specimens to be used for DNA, RNA, and protein extraction were stored as fresh frozen samples. Usage of specimens for research/ within this project are under approval from the Ethical Review Board (Etikprövningsnämnden) with number DNR 2010/127.

**Table 1.**
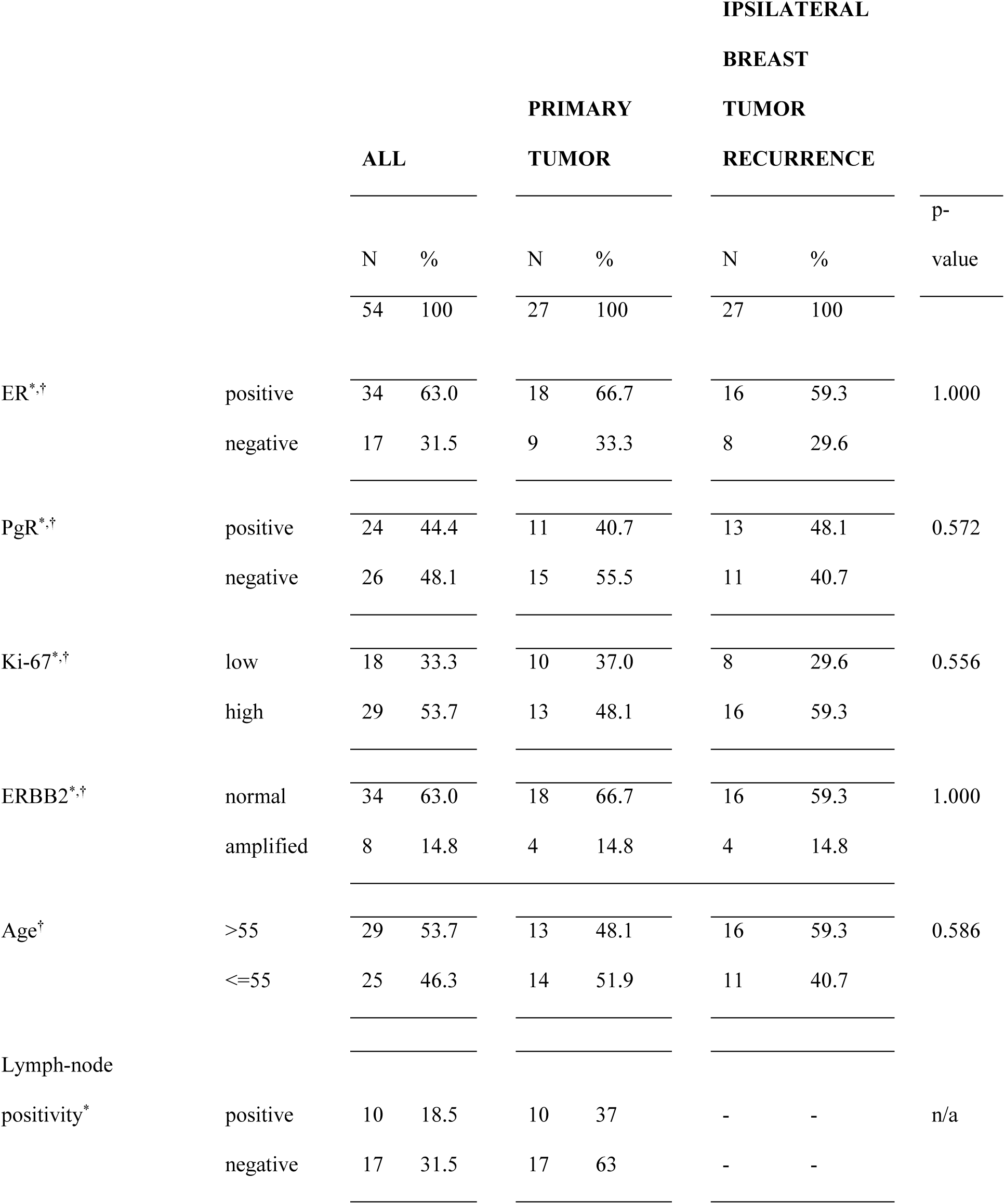
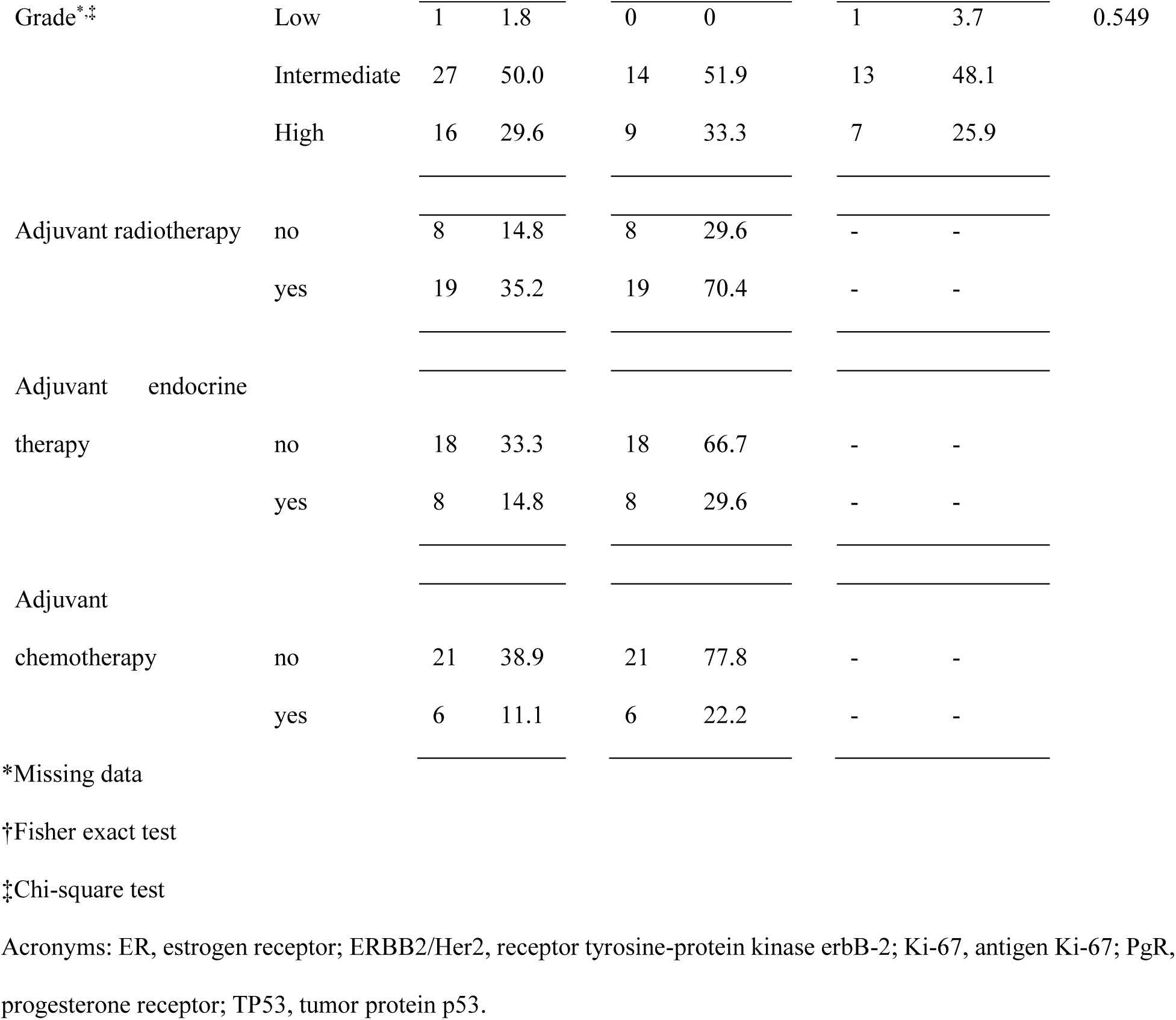
Clinical variables.

### Immunohistochemical routine biomarker analysis

FFPE tissues were cut into 3-4 µm sections and put on TOMO slides (MG-TOM-11/90, Histolab). Evaluation of ER (immunohistochemistry, IHC; staining cutoff: 10% positive cells), PgR (IHC; staining cutoff: 10% positive cells), ERBB2/Her2 (IHC and *in situ* hybridization for equivocal cases) and Ki-67 (staining cutoff: 30% positive cells) were performed according to routine clinical practice in Sweden. Briefly, antibodies used for IHC stainings were: ER: clone SP1, 790-4324 Ventana; PgR: clone 1E2, 790-2223 Ventana; Ki-67: clone MIB-1, M7240 DAKO (dilution 1:100); HER2: clone 4B5, 790-2991. Slides for ER, PgR, and Ki-67 evaluation were stained on the Discovery ULTRA (Ventana Medical System Inc., Tucson, AZ, USA). HER2 staining was performed on the Benchmark ULTRA (Ventana Medical System Inc., Tucson, AZ, USA). For all IHC, ULTRA cell conditioning (ULTRA CC1) pH 8-8.5, was used for heat-induced epitope retrieval. The primary antibodies were incubated for 32min and visualized with conventional 3,3’-diaminobenzidine IHC detection kit.

### DNA, RNA, and protein extraction

Breast tumor tissues (PTs and IBTRs) were processed using the AllPrep DNA/RNA/Protein (Qiagen) protocol. Tissue lysis was performed by re-suspending ∼30mg of sliced frozen tissue in a solution containing 1% β-mercaptoethanol in RLT buffer (supplemented with antifoam agent; ID 19088, Qiagen). Next, steel beads (ID 79656, Qiagen) were added and samples were incubated in a Tissue Lyser LT (Qiagen) for 4min at 50Hz. Steel beads were then removed and 400µL of 1% β-mercaptoethanol in RLT buffer was added to samples, which were then centrifuged at 14,000xg for 5min. Supernatants were transferred to new tubes, and then frozen at -80°C. DNA, RNA, and protein extraction were performed according to manufacturer instructions (AllPrep DNA/RNA/Protein minikit; Qiagen). Each spin column flowthrough (DNA, RNA, protein) was stored at -80°C until analysis (sequencing or mass spectrometry; MS).

### Whole-genome sequencing

Sample library was performed twice for every sample, using a PCR-free method for specimens with high DNA yield, and employing a PCR amplification step for low yield samples. PCR-free libraries were prepared from 1μg DNA using the TruSeq PCRfree DNA sample preparation kit (cat# FC-121-3001/3002, Illumina) targeting an insert size of 350bp. PCR-amplified sequencing libraries were prepared from 100ng DNA using the TruSeq Nano DNA sample preparation kit (cat# FC-121-4001/4002, Illumina) targeting an insert size of 350bp. Both library preparations were performed according to manufacturers’ instructions. Paired-end DNA sequencing with 150bp read length was performed at the SNP&SEQ Technology Platform in Uppsala (Uppsala University, Uppsala, Sweden) using an Illumina HiSeqX sequencer (Illumina, San Diego, CA) with v2.5 sequencing chemistry.

### Variant calling

Alignment to reference genome GRCh38 was performed using bwa’s (v0.7.13) BWA-MEM algorithm, and conversion to BAM format and coordinate sorting was performed using samtools (v1.3). Duplicates were marked using Picard (v2.0.1). To identify all possible active regions and ensure that all samples had their information represented comparably the tools RealignerTargetCreator and IndelRealigner from GATK (v3.7) were used. Samples were processed with a scatter-gather methodology, dividing each sample by chromosome to identify and realign any misaligned reads in active regions. Samples were then merged using Picard MergeSamFiles. GATK 3.7 BaseRecalibrator and PrintReads were used to identify potential systematic errors in the data and recalibrate the base quality scores.

A panel of normal (PoN) variants was created and used as a blacklist during variant calling. First, variants were called using GATK (v3.8) MuTect2 with only a normal sample as input and then CombineVariants to aggregate the output for all normal samples. Only variants observed in at least two samples were included. For further variant calling, pairs of matched normal and tumor samples (2 out of 27 patients: S12 and S18) were called together if the patient had a matched normal sample, otherwise the tumor sample was called alone. In either case the PoN was used. A scatter-gather methodology was used to optimize runtimes, and CatVariants was used to merge the variants. The variants were filtered using the built in Mutect2 filtering (cutoff: lack of PASS annotation). The variants were annotated using snpEff^27^ (v4.2) and annovar (v2017.07.16). Information on the allele frequency of variants in population databases SweGen, ExAC and gnomAD was also added together with COSMIC database annotation. For the SweGen and ExAC database annotation, a lift-over of the variant files was performed using Picard (v2.10.3) with the LiftoverVcf command to obtain GRCh38 coordinates.

We applied the TPES^28^ (v1.0.0; https://cran.r-project.org/web/packages/TPES/index.html) package to estimate tumor purity values for all cancer samples using the single nucleotide variant (SNV) list as input. These estimates were used to filter the SNVs.

The filters are applied in order as follows:

1. Population allelic frequency (AF) filter: the observed AF in gnomAD and SweFreq needed to be 0 or NA (i.e. variant has never been observed in a sample from these datasets).
2. Allelic Depth (Support) filter: the number of reads supporting the variant in the tumor needed to be larger or equal to 2. Additionally, if a matched normal was available, the number of reads supporting the variant in that sample needed to be 1 or 0.
3. Coverage filter: the total number of reads overlapping the position needed to be 10 or more in the tumor sample. If a matched normal sample was available, that needed to have 10 or more reads coverage.
4. PoN filter: variant not present in the PoN file (i.e. it cannot have been detected in any of the normal samples from this dataset).
5. TPES filter: the Log2 ratio of the probability of a given variant being observed under the cancer vs. background model needed to be ≤ –1 (i.e. removal of variants where the background model is twice as likely to produce the observed variant). Specifics on calculations are presented here:

a. For each sample *i* with a TPES^28^ based tumor purity estimate (*t_i_*) we define 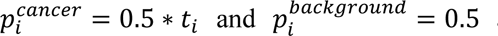 as the probabilities of a binomial distribution for the heterozygous SNV model, analogous with 1.0 instead of 0.5 for the homozygous SNV model. If no TPES based tumor purity estimate *(t_i_)* existed for a given sample *i*, this filtering step was skipped
b. We then calculate for each variant *j* the ratio 2 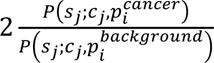, where *s_j_* is the support of variant *j* in sample *i* and *c_j_* is the coverage of that variant. Here, *P(k;n;p)* is the probability of observing exactly *k* out of *n* hits (i.e. reads with alternative allele) under a binomial distribution with probability *p*.
c. If the Log2 ratio is larger (>) than -1 for the heterozygous or the homozygous case, the variant is kept. If it is below or at -1 in (≤) both in the homo- and heterozygous cases, indicating a higher probability in the background model, the variant is filtered out.

The SNV list was derived by extracting the mutations contained in the union of the COSMIC cancer gene census^29^, the FoundationOne® gene list^30^, genes part of the Memorial Sloan Kettering IMPACT platform^31^, and the list of reported BC driver genes^12^. Variants were filtered based on impact (moderate or high were included) and type of variant (downstream gene variant, upstream gene variant, 3’ UTR variant, 5’ UTR variant, and synonymous variant cases were excluded). Mutational signatures were determined using the MutationalPatterns package (v3.3.0)^32^ by fitting the SNV counts per 96 tri-nucleotide context to the 30 COSMIC signatures^29^.

We used sciClone (v1.1.0; https://github.com/genome/sciclone) to build the clustering of SNVs by their variant allele frequencies. Applying clonevol (v0.99.11; https://github.com/hdng/clonevol) to sciClone-derived clustering did not yield any valid model of tumor evolution.

### Copy Number call

CN calls were obtained by determining total coverage across the genome in 10kb bins for each sample, then using the R locfit.robust function to fit the relationship between GC content and bin coverage, then adjusting for the differences in GC-coverage relationships across samples. The resulting adjusted coverage values were converted into Log2 ratios by employing either the matched normal sample (if available), or the median adjusted coverage of the sample itself as denominator. The Log2 ratios were then centered (median subtraction), adjusting for between-sample coverage differences.

CN segmentation was performed on the centered Log2 ratios using the circular binary segmentation algorithm implemented in the DNAcopy R/Bioconductor package. The resulting CN segments were then mapped to genes by finding overlaps with annotated exons of each gene. For genes overlapping multiple copy number segments the CN values were averaged.

To determine CN gains and losses between paired PTs and IBTRs a CN delta was calculated with the following formula:

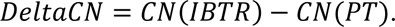

CN changes were taken into account only if they impacted genes that showed a minimum CN of 0.5 in matched PT and IBTR samples and where DeltaCN was above 0.75 (gain) or below -0.75 (loss).

### RNA sequencing

RNAseq was performed as previously reported^21^. Briefly, the amount, concentration and quality of the extracted RNA was tested using a Bioanalyzer 2100 instrument (Agilent Technologies), a NanoDrop ND-1000 spectrophotometer (Thermo Fisher Scientific) or Caliper HT RNA LabChip (Perkin Elmer). All samples had a RNA integrity value of 6.0 or higher.

RNAseq library preparation and analysis were conducted as previously described^33^. Briefly, 100 ng of RNA input was used for cDNA library preparation using the TruSeq® Stranded mRNA NeoPrep kit (Illumina), according to manufacturer instructions. Concentration of cDNA was measured (QuantIT® dsDNA HS Assay Kit; Thermo-Fisher), and libraries were then denatured and diluted according to the NextSeq® 500 System Guide (Illumina). RNAseq was then performed on a NextSeq 500 (Illumina) sequencer generating paired-end reads of length 75bp.

### RNAseq data processing

De-multiplexed RNA-Seq reads were aligned to the GRCh38 human reference genome using STAR aligner (v020201) with an overhang value of 75 to match the read-length. The standard GATK analysis pipeline was then applied (GATK; v3.7-0-gcfedb67). The resulting alignment files were processed by first generating per-gene read counts mapping to the GRCh38 GTF file from Ensembl (v95) using the *summarizeOverlaps* function in “Union” mode to count reads that uniquely mapping to exactly one exon of a gene (GenomicAligner, v1.18.1). Next, genes with no counts in any of the samples were discarded, and DESeq2 analyses using R/Bioconductor package (v1.22.2) were performed^34^.

### Protein digestion

Protein flow-throughs from the AllPrep protocol were precipitated in ice-cold (-20°C) methanol, as previously described^35^. Briefly, protein pellets were then suspended in 100mM Tris (pH 8.0) buffer containing 100mM dithiothreitol and 4% w/V sodium-dodecyl-sulphate and incubated at 95°C for 30min under mild agitation. Samples were then cooled to room temperature, diluted in 8 M urea in 100mM Tris (pH 8.0) buffer, loaded on 30KDa molecular filters (Millipore) and centrifuged at 14,000xg for 20min. Filters were washed with urea buffer and centrifuged at 14,000xg for 10min. Proteins were alkylated with iodoacetamide in urea buffer (30min in the dark), washed with urea buffer and tri-ethyl-ammonium bicarbonate buffer (pH 8.0), and trypsin was added (enzyme-protein ratio 1:50; incubation at 37°C for 16h, 600RPM). Filters were then centrifuged at 14,000xg for 20min to retrieve tryptic peptides, loaded onto C18 (3 stacked layers; 66883-U, Sigma) stage tips (pretreated with methanol, 0.1% formic acid (FA) in 80% acetonitrile solution, and 0.1% FA in ultrapure water), washed with 0.1% FA in ultrapure water solution, and eluted with 0.1% FA in 80% acetonitrile. Eluates were then dried and subjected to SP3 peptide purification, as previously described^36^. Briefly, 2µL of SP3 beads (1:1 ratio of Sera Mag A and Sera Mag B re-suspended in ultrapure water; Sigma) were added to dried peptides and incubated for 2min under gentle agitation. A volume of 200µL of acetonitrile was then added and samples were incubated for 10min under agitation. Sample vials were then placed on a magnetic rack and washed again with acetonitrile for 10min. Elution was performed by adding 200µL of 2% dimethyl sulfoxide in ultrapure water to the bead-peptide mixtures and incubating them for 5min under agitation. Supernatants were then collected, dried, and stored at -80°C until MS analysis.

### Mass spectrometry analysis

Tryptic peptide mixtures were subjected to data-independent acquisition (DIA) MS analysis. Samples were eluted in a 120min gradient (flow: 300 nl/min; mobile phase A: 0.1% FA in ultrapure water; mobile phase B: 80% acetonitrile and 0.1% FA) on a Q-Exactive HFX (Thermo-Fisher) instrument coupled online to an EASY-nLC 1200 system (Thermo-Fisher). Digested peptides were separated by reverse phase HPLC (ID 75µm × 50cm C18 2µm 100Å resin; Thermo-Fisher). Gradient was run as follows: 10-30% B in 90min; 30-45% B in 20min; 45-90% B in 30s, and 90% B for 9min. One high resolution MS scan (resolution: 60,000 at 200m/z) was performed and followed by a set of 32 DIA MS cycles with variable isolation windows (resolution: 30,000 at 200m/z; isolation windows: 13, 14, 15, 16, 17, 18, 20, 22, 23, 25, 29, 37, 45, 51, 66, 132m/z; overlap between windows: 0.5m/z). Ions within each window were fragmented by HCD (collision energy: 30). Automatic gain control target was set to 1e6 for both MS and MS/MS scans, with ion accumulation time set to 100ms and 120ms for MS and MS/MS, respectively. Protein intensities were derived by employing our previously established computational workflow^21^. A total of 4,640 proteins were identified after FDR filtering (cutoff: 0.01). Batch effect correction was performed using the limma^37^ (v3.46.0) package. Raw protein intensities were Log2 transformed and centered prior differential expression analysis by DEqMS^38^ (v1.8.0).

### Statistical and pathway analyses

All statistical tests were performed in R (v4.0.5; correlations, hierarchical clustering, and differential expression tests) or GraphPAD (v.9; contingency tables for Fisher and chi-square tests).

Gene Set Enrichment Analysis (GSEA; v4.1.0)^39^ was performed on scaled and Log2 transformed RNA and protein tables. Databases: Hallmarks (v5.2), ALL (v5.2); permutation type: gene set; scoring: classic; metric: t test; other parameters were kept at default settings; significance cutoff: FDR < 0.25.

## Results

### Proteogenomic validation of clinical and molecular characteristics

In this study we analyzed a set of 54 samples from 27 patients who developed IBTR. The 27 tumor pairs (PTs and IBTRs) were selected from a previous large multi-center study that aimed to define radiosensitivity markers^26^. The paired analysis of PT and IBTR enabled a patient-centered view of tumor evolution, measured by the changes in the genomic, transcriptomic, and proteomic landscape. Both PT and IBTR were stratified based on ER-, PgR-, ERBB2-, and Ki-67-status, histological tumor grade, molecular subtype^40^ and treatments (**Figure 1A-B**). No statistical differences in clinical and histopathological parameters were observed between PTs and IBTRs (**Table 1** and **Table S1**). IHC analysis of the ER and PgR revealed no significant differences for the tumor proliferation marker Ki-67 in the primary tumors, corroborated by transcriptomic and proteomic measurements (**Figure 1C-D**). In contrast, there was a significant difference of Ki-67 in IBTRs (RNA p < 0.01, protein p = 0.019). This observation may stem from heterogeneous Ki-67 expression in PT and IBTRs due to clonal selection. Interestingly, 10 tumor pairs switched tumor marker status in the transition from PT to IBTR (ER: n = 1; PgR: n = 6; Ki-67: n = 7), which were in most cases validated by proteogenomics (RNA level: ER n = 1/1, PgR n = 4/6, Ki-67 n = 1/7; protein level: PgR n = 4/6, Ki-67 n = 1/7; **Figure S1A-B**). The weak correlation of Ki-67 status switch to transcript and protein levels might be due to the discrepancy between transcript and protein Ki-67 measurements when compared to immunohistochemistry, which itself depends on both analytical and pre-analytical factors^41, 42^. ERBB2/Her2 status was confirmed at the CN, RNA, and protein level (**Figure 1E**), with no status switch between PT-IBTR pairs. Overall, these results demonstrate concordance between biomarker status and the techniques employed in this study as well as pinpointing relevant changes in clinical tumor markers between PTs and IBTRs.

**Figure 1.**
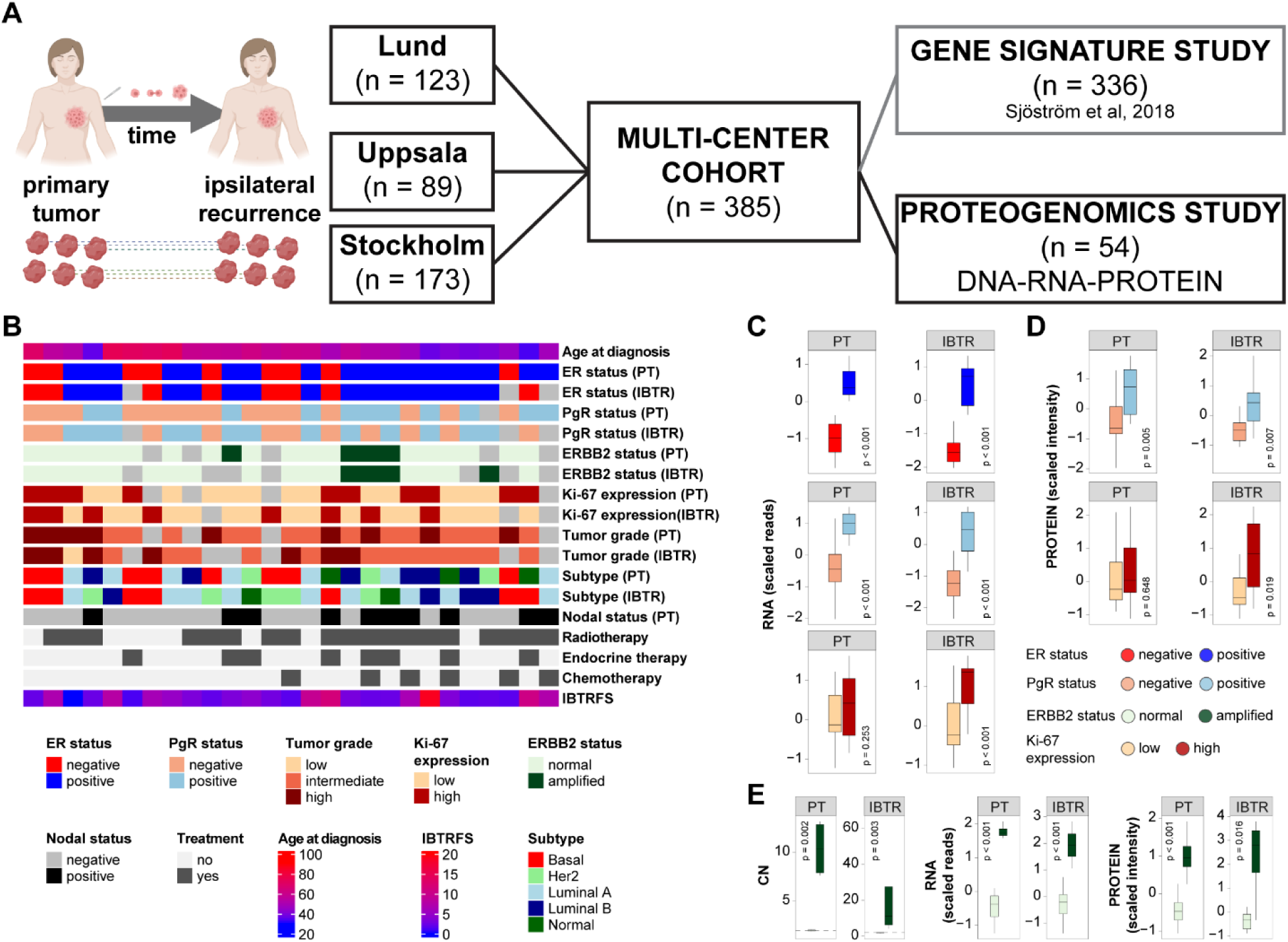
Cohort selection and metadata. We selected a set of PTs and matched IBTRs (n = 27 + 27) from a larger multi-center (Lund, Uppsala, Stockholm) study (Panel A). Clinical data and histopathological characteristics were registered upon sample collection or after analyses of paraffin-embedded material, if available. Panel B: Description of key clinical parameters (light gray boxes represent missing values). Frozen tumors were analyzed by WGS, RNAseq, and proteomics. Panel C: Estrogen and Progesterone receptor levels measured by RNA. Panel D: PgR and Ki-67 status levels validated at the protein level. Panel E: Comparison between pathological analysis and CN, transcript, and protein levels of ERBB2/Her2. Acronyms: CN, copy number; ER, estrogen receptor; ERBB2/Her2, receptor tyrosine-protein kinase erbB-2; Ki-67, antigen Ki-67; IBTR, ipsilateral breast tumor recurrence; IBTRFS, IBTR-free survival; PgR, progesterone receptor; PT, primary tumor.

### Changes in mutational signatures between primary and recurrent tumors

Mutational processes involved in breast cancer recurrence are in many cases a result of homologous recombination deficiency, APOBEC-mediated mutagenesis, or age-related genome deterioration^12, 20, 43^. To quantify the magnitude of genomic changes between matched PTs and IBTRs, we first analyzed the frequency of base transitions and transversions. In this analysis, the contribution of 30 previously published mutational signatures^18^ were determined in our dataset (**Figure S2A**). Here, two signatures were detected at a high frequency across all samples (referred to as high contribution), with signature 3, enriched in cytosine transversions (possible cause: failure of DNA double-strand break repair by homologous recombination), and signature 5, enriched in cytosine and thymine substitutions (possible cause: unknown), as the most contributing signatures in both primary and recurrent tumors (**Figure S2B**). Apart from signature 3, a low contribution was observed for signatures previously associated to BC i.e. signature 8, 13, 17, 18^19^. Association analysis between mutational signatures 3 and 5 and the clinical and histopathological features revealed a significant relationship between loss of ER expression and signature 3. As signature 3 has been associated to deficient DNA repair during replication, these results suggest a link between the establishment of this mutational mechanism and absence of ER. In addition, there was a significantly higher contribution of signature 1 in tumors that express ER and *wt TP53*. This indicates that the mutational burden, and its changes during tumor progression and evolution, may be modulated by either TP53 independent factors or non-genomic mechanisms, such as proliferation rate (**Figure S2C**).

In the next step, we compared changes in contribution of the molecular signatures between matched PTs and IBTRs (**Figure 2A**). This analysis showed that the contribution of signature 3 was increased in IBTRs, while the contribution of signature 5 was decreased (**Figure 2B-C**). Interestingly, the increased contribution of signature 3 was significantly associated with absence of hormonal receptor (Wilcoxon test, ER p = 0.010; PgR p = 0.003), high proliferation rates (Ki-67 p = 0.025, tumor grade p = 0.007), and radiotherapy treatment (p = 0.029; **Figure 2D**). As signature 3 has been associated with failure of double strand break repair by homologous recombination, such a process might be exacerbated in tumors with high proliferation rates (hormonal receptor negative, high Ki-67, and high grade tumors). On top of this, the DNA damage caused by radiotherapy might have further enhanced this process.

**Figure 2.**
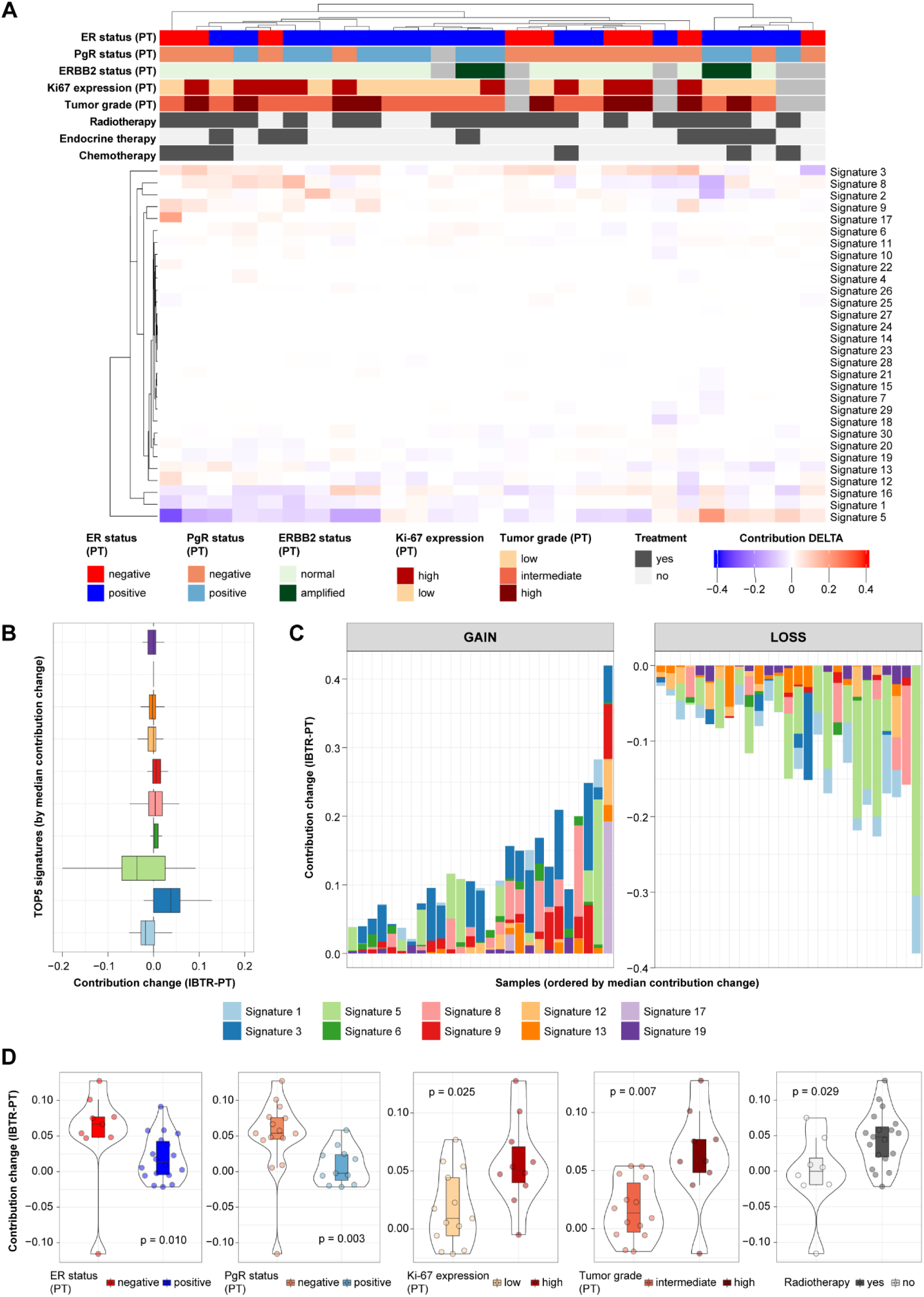
Mutational spectrum and shift of mutational signature contributions. We evaluated the contribution of the 30 COSMIC mutational signatures within our samples (PT and IBTR subsets). Contribution delta was then calculated as a measure of mutational process evolution for each PT-IBTR pair. Panel A displays hierarchical clustering of mutational contribution delta (IBTR-PT; light gray boxes represent missing values). Panel B shows the top 10 signatures with contribution changes between primary and locally recurrent tumor pairs. Bar charts displaying the evolution of (top 10, from panel A) mutational signatures between primary and locally recurrent tumors are shown in panel C. Significant associations between changes in mutational signature 3 contribution and clinical variables are depicted in panel D. Acronyms: ER, estrogen receptor; ERBB2/Her2, receptor tyrosine-protein kinase erbB-2; Ki-67, antigen Ki-67; IBTR, ipsilateral breast tumor recurrence; IBTRFS, IBTR-free survival; PgR, progesterone receptor; PT, primary tumor.

In contrast, the reduction in contribution of signature 5 was significantly associated with ERBB2 status (p = 0.026; **Figure S3**), suggesting a relationship between the mutational processes underlying this signature and the enhanced kinase activity of ERBB2-amplified tumors. Signature 9, which has previously been associated to POLH-mediated mutagenesis, showed a significant association with radiotherapy (p = 0.011) and a significant negative correlation with age (Rho = - 0.485; **Figure S3**). Signature 12, similarly to signature 3, significantly increased in contribution in ER negative PT-IBTR pairs (p = 0.047), while the APOBEC associated signature 13 showed increased in contribution in tumors treated with radiotherapy (p = 0.034; **Figure S3**), suggesting a possible role of APOBEC in radiotherapy resistance. Other signatures were found significantly associated with clinical biomarkers, but their contribution in the dataset was much lower (i.e. below 0.01) and were deemed as minor factors for changes in the mutational landscape in our samples. In conclusion, these results show that the spectrum of mutational signatures changes between PTs and IBTRs, and that key tumor features such as presence/absence of hormonal receptors or ERBB2 amplification influence the impact of mutational processes during tumor evolution.

### Copy number and mutational changes in ipsilateral breast tumor recurrences

As we observed a switch in tumor markers and a significant change in the contribution of two mutational signatures between matched PTs and IBTRs, we hypothesized that these events were accompanied by additional genomic changes. To address this, we analyzed the frequency of CN alterations and single nucleotide variants. We calculated genome-wide CN changes (DeltaCN, see *Methods*) between PT-IBTR pairs per chromosome and characterized as either gains or losses (cutoff DeltaCN: ±0.75; median gain/sample: 363, IQR: 48.5-1115.5.; median loss/sample: 95, IQR: 22-291; **Figure S4A**). Closer inspection of the top 10 CN gain and losses in each chromosome revealed that genomic regions in chromosomes 8 and 17 were frequently amplified or deleted in our sample set (**Figure S4B-C**). Overall, we did not detect any association between changes in CN in the PT-IBTR pairs and CN occurrence at specific chromosomes. Interestingly however, clustering of the CN changes showed a relationship between the frequency of gain/loss and hormonal receptor status (**Figure 3A**). Association analysis to clinical biomarkers confirmed that loss of ER and PgR as well as high Ki-67 expression were all associated with a significant increase in gene gains (Wilcoxon test, ER p = 0.059, PgR p = 0.017, Ki-67 p = 0.005; **Figure 3B**). These results suggest an association between CN gains and the absence of the ER-mediated transcriptional program as well as high proliferation rates. To search for molecular drivers of these relationships, we investigated whether the frequency of CN gain was linked to the expression of mutated *TP53*, which typically leads to genomic instability^44^. Despite the fact that *TP53* mutations are frequent in ER negative BCs, as also observed in our dataset (Fisher p = 0.029; **Figure 3A**), no significant association between CN gains and *TP53* mutational status was observed (p = 0.099; **Figure S5**), suggesting other factors play a role in the establishment of CN changes in this sample set.

**Figure 3.**
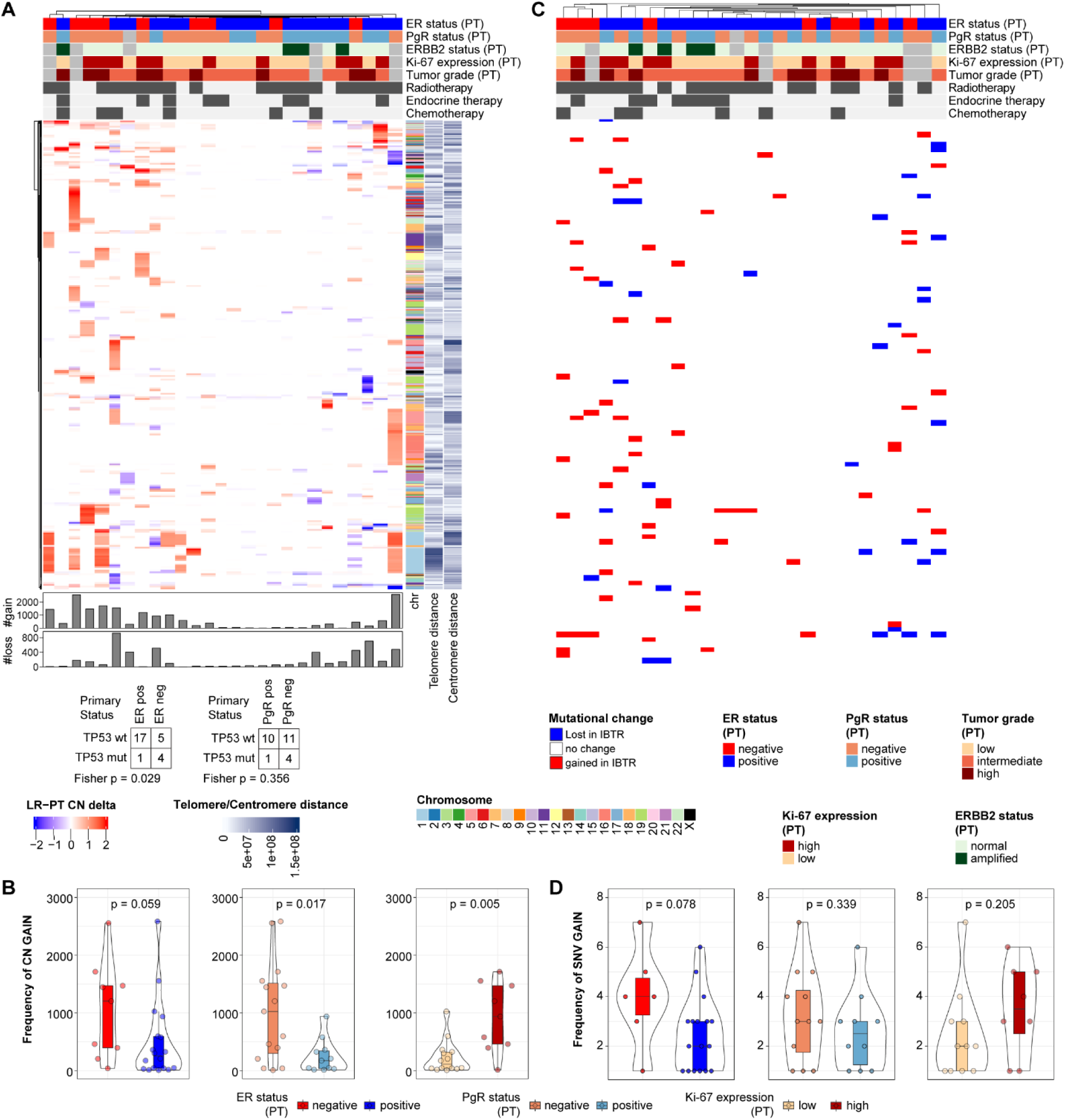
Changes in copy number and key drivers. Copy number changes between paired tumors and mutational (SNV) status of COSMIC cancer genes was evaluated in our cohort. Panel A: Heatmap of copy number changes between primary and recurrent tumors. Bottom bar charts display sample-wise frequencies of IBTR CN gain and loss over the matched PT. Panel B: Association of copy number gain between primary and recurrent tumors to key biomarkers. Panel C: Heatmap of mutational status change (i.e. gain or loss in IBTR) of cancer genes from COSMIC. Panels D: Association of mutational gain between primary and recurrent tumors to key biomarkers. Light gray boxes in heatmaps represent missing values. Acronyms: CN, copy number; ER, estrogen receptor; Ki-67, antigen Ki-67; IBTR, ipsilateral breast tumor recurrence; IBTRFS, IBTR-free survival; PgR, progesterone receptor; PT, primary tumor; SNV, single nucleotide variant.

We then tested whether there was a relationship between CN gains and adjuvant therapy (endocrine, chemotherapy, and radiotherapy), though no significant association was found. Furthermore, we observed no correlation between CN gains and age at diagnosis (of PT), and only a weak correlation with IBTRFS (Spearman Rho = 0.314; **Figure S5**).

Next, we analyzed SNV changes in the for a set of previously defined key cancer genes (Nik-Zainal et al^12^; **Figure S6**). Here we evaluated the SNV gain/loss occurring in PT-IBTR pairs. This analysis showed that the most common SNV changes with medium or high impact were missense and stop codon gains (**Figure S7A**), with a general trend towards an increasing SNV burden in IBTR (**Figure S7B**). Further analysis showed that ER negative tumors increase in SNV gains (Wilcoxon p = 0.078; **Figure 3C-D**), while no significant association was observed between SNV gains and other biomarkers or clinical features (**Figure 3D** and **Figure S8**) Upon assessing the most frequently mutated genes within ER positive and negative tumors, we confirmed that *PIK3CA* and *TP53* were the most (commonly) mutated genes in these subgroups, respectively (**Figure S7C**). These mutations were largely maintained or expanded through clonal selection in IBTRs possibly due to a conferred selective advantage towards cancer growth and survival.

Alongside CN and SNV gains, which in turn constitute a measure of tumor genomic evolution and/or clonal expansion from PTs to IBTRs, we detected several losses: median SNV gain/sample = 3 (IQR: 1-4), median SNV loss/sample = 2 (IQR: 1.25-3). These likely indicate a reduction or loss of tumor sub-clones during from PT to IBTR, but did not associate with hormonal receptor status or other clinical variables with the exception of weak positive relationships with age at diagnosis of PT (Spearman Rho = 0.295) and IBTRFS (Spearman Rho = 0.372; **Figure S9**). Taken together, the paired analysis conducted here shows that primary ER and PgR negative tumors were more genomically unstable (as also reviewed in ^45^), displaying a higher tendency to acquire genomic changes such as CN and SNV, resulting in highly mutated IBTRs.

### Multi-omic evolution of primary breast cancer

Having established that the absence of ER is significantly associated with the accumulation of CN and SNV in locally recurrent tumors in comparison with their primary counterparts, we investigated to what degree the genomic changes translated into alterations at the transcriptome and proteome levels.

With this in mind, we calculated Euclidean distances between each sample within our genomic (CN and SNV), transcriptomic, and proteomic datasets (**Figure S10**). Next, we extracted distances between PT-IBTR pairs. A higher distance indicated highly different IBTRs when compared to their matched PTs. As a major factor in determining accurate genomic (CN, SNV) and molecular (RNA, protein) measurements^28^, tumor purity was ruled out as a potential confounder of PT-IBTR distance across all omics levels (**Figure S11**).

We observed that distances between PT-IBTR pairs in general were greater at the RNA and protein levels when compared to CN and SNV (**Figure S12A**), indicating that genomic changes during tumor evolution either have larger repercussion at the expression level, or that other non-genomic mechanisms contribute to modulate the transcriptome and proteome^46^. Upon assessing the distribution of PT-IBTR distances we noticed similarities between CN, SNV, and RNA distance metrics, but much less for protein-wise distances, suggesting that changes at the genomic level are have a greater impact at the transcript level, but to a much lesser degree at the protein level (**Figure S12B**). Correlation analysis between PT-IBTR distances showed a positive association between all four *omics* levels (**Figure S12C**). Spearman correlations coefficients showed a strong correlation between CN- and RNA-distances (Spearman Rho = 0.700), while protein- and SNV-distances clustered separately. The co-clustering of CN and RNA-based distances is likely due to a closer relationship between gene CN and transcript levels, while SNV and protein distances clustered separately as they may impact activity rather than expression levels (SNV), or are under different regulatory mechanisms (protein; e.g. transcription rate *vs* translation rate), respectively. Next, we assessed whether each PT primarily associated with its matching IBTR by displaying if the minimum PT distance was linked to its matching IBTR or another sample. Hierarchical clustering analysis of sample-wise CN, SNV, RNA, and protein level distances (**Figure 4A**) showed that only 3 PTs had their respective IBTR as closest neighbor across all data layers, suggesting that changes in genomic features directly alter transcript and protein expression only in a small subset of tumors. As new PTs may be misdiagnosed as IBTRs of previous malignancies^47, 48^, we compared the clonal evolution of PT-IBTR pairs. Here, sample pairs with a matched normal showed an overlap between variant allele frequencies (**Figure S13**), thus we concluded that IBTRs derived from their respective PTs in these patients.

**Figure 4.**
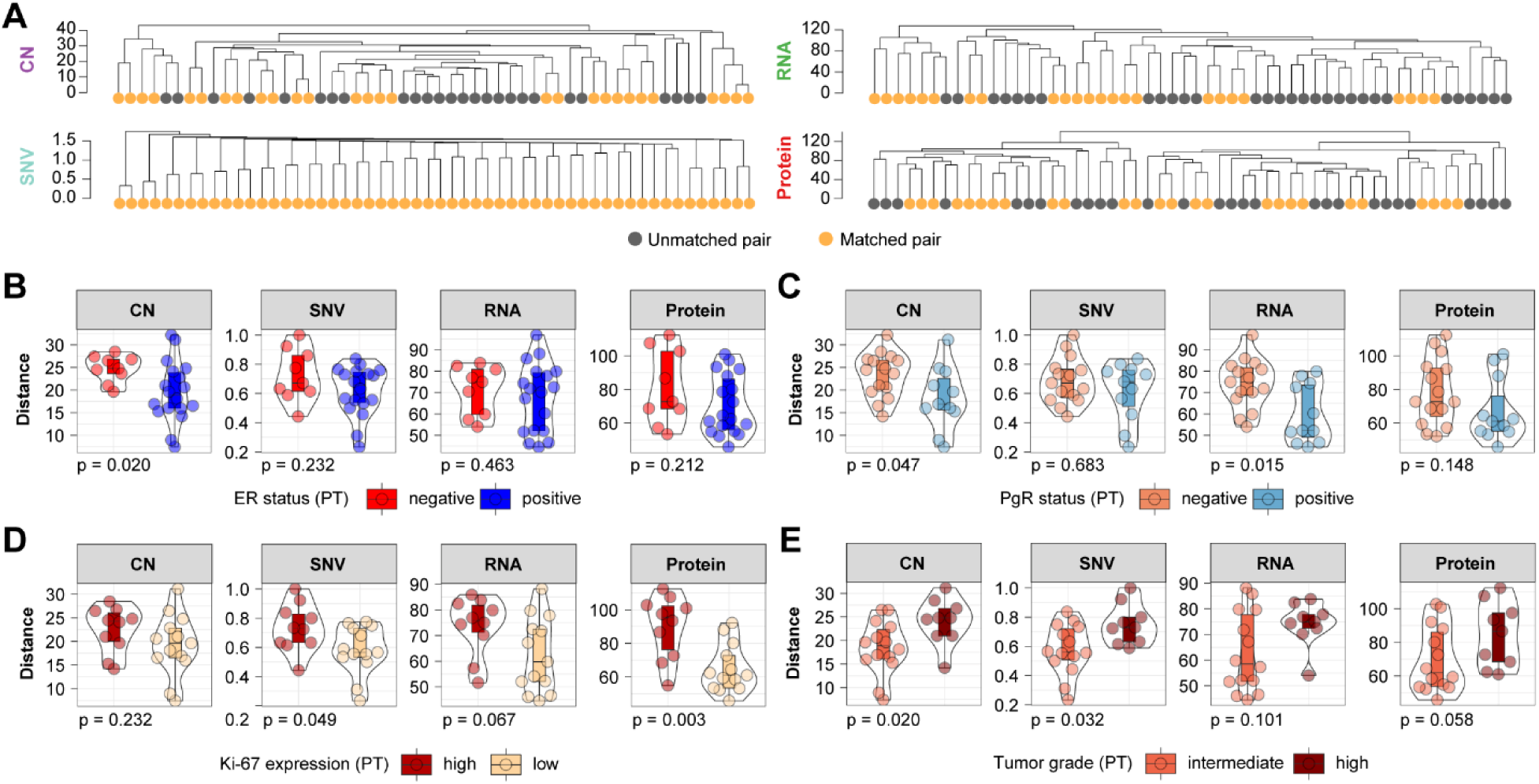
Multi-omic drift assessment of breast cancer recurrences. Euclidean distances were calculated between tumor pairs at the CN, SNV, transcript, and protein levels as a measure of evolutionary drift. Panel A: PT-IBTR distance-based clustering at the CN, SNV, transcriptome, and protein levels. Panels B-E: Association between clinical variables and PT-IBTR distances across data layers. Acronyms: CN, copy number; ER, estrogen receptor; Ki-67, antigen Ki-67; IBTR, ipsilateral breast tumor recurrence; PgR, progesterone receptor; PT, primary tumor; SNV, single nucleotide variant; TP53, tumor protein p53.

Overall, with the exception of the SNV layer where every PT matched with its correspondent IBTR sample, we observed that the matching of PT-IBTR pairs varied in relation to data layer indicating that the changes between each tumor pair are dependent on different mechanisms, such as promoter methylation, histone binding, kinase activation, microenvironment signaling could play a key role in defining tumor evolution.

To determine factors associated with IBTR evolution, we assessed the relationship between distances and clinical and histo-pathological features of the cohort. Here, a strong inverse relationship was observed between distances and shared PT-IBTR SNVs (Spearman Rho range: - 0.399 to -0.952; **Figure S14A**), indicating that a lower level of shared mutations between PT and IBTR is related to larger distances and more dissimilar PT-IBTR pairs, which is turn is reflected in more dissimilar gene expression and protein abundance patterns. Overall, these results indicate that mutational drift is established together with CN changes, and it is reflected by changes in gene and protein expression.

As with our previous observations when assessing changes in mutational signatures (**Figure 2**), CN, and SNV (**Figure 3**), we argued that clinical and histopathological characteristics of each PT might be determinants of its evolution into an IBTR. Association testing revealed a higher mutational, CN, and transcript/protein expression distance between tumor pairs in hormonal receptor negative, Ki-67 high, and high grade cancers (**Figure 4B-E**). These results confirm our previous analyses (**Figure 2-3**), and indicate that more substantial changes at the genomic, transcriptomic, and proteomic levels between PT-IBTR pairs might directly stem from high proliferative activity and other features typical of ER/PgR negative cancers. With the exception of hormonal receptor status and Ki-67 levels, no significant association was observed for other clinical or histopathological variables (**Figure S14-15**). These results confirm that hormonal receptor negative and high proliferating PT tumors often result in IBTR with high frequency of CN and SNV changes which in turn promote transcriptome and proteome reprogramming.

### Differential transcriptome and proteome evolution of ER positive and ER negative tumors

We observed that lack of hormonal receptor and elevated Ki-67 levels in primary tumors are associated to genomic instability that affect both the CN and mutational landscapes, we evaluated to what degree ER status impacted the transcriptome and the proteome in this sample cohort. Differential expression analysis between ER positive and ER negative PT-IBTR pairs showed a set of overlapping transcripts and proteins (**Figure 5A-B**). These transcript and protein pairs were enriched for mTOR signaling (e.g. mTORC1 signaling) and immune response (e.g. allograft rejection) pathways enriched in IBTRs and PTs, respectively (**Figure 5C-F**). The evolution of PTs into IBTRs and the consequent expansion of the mutational landscape could explain the dysregulation of proliferation-related pathways such as mTOR. In contrast, the enrichment of inflammation and immune system-related signaling in PTs might indicate changes in the relationship between the cancer and its microenvironment, possibly geared towards immune evasion. Closer inspection of these pathways revealed enhanced expression of inflammatory cytokines such as IL6 and IL8 as well as matrix remodeling enzymes (MMP9; **Figure S16A-B**), indicating a tendency towards invasion of the surrounding tissue.

**Figure 5.**
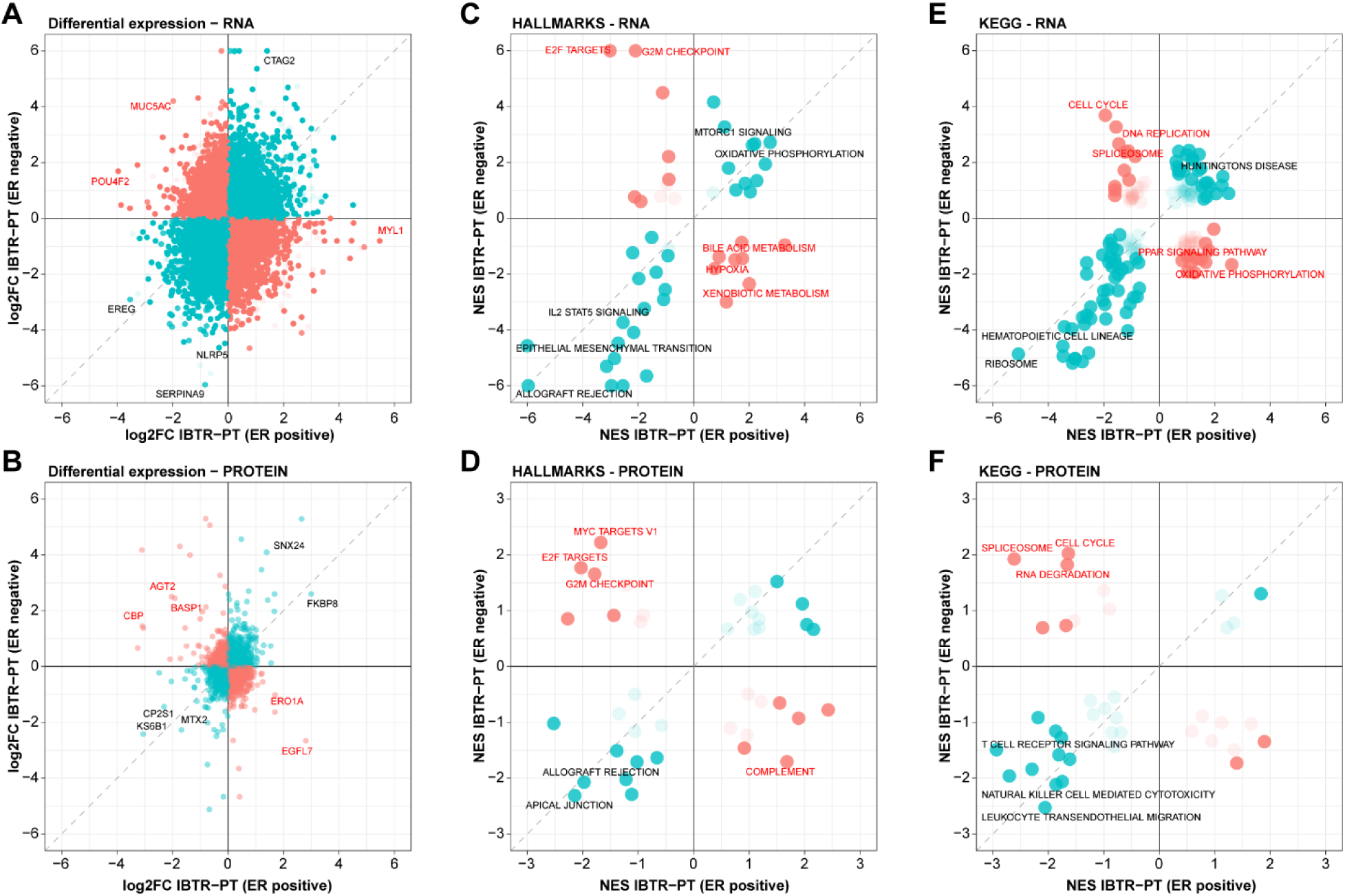
Estrogen receptor expression-dependent evolution of recurrent breast cancers. Differential expression and pathway enrichment analyses were performed between paired IBTR and PT specimen within the ER positive and ER negative groups. Results were compared to measure the degree of deviation in the evolution of ER positive and ER negative tumors. Panel A-B: Differential transcript (A) and protein (B) levels between PT and IBTR samples in ER positive and negative tumors. Panel C-F: Pathway enrichment divergence in ER positive and negative patients at the RNA and protein levels. Acronyms: ER, estrogen receptor; IBTR, ipsilateral breast tumor recurrence; PT, primary tumor.

Genes and pathways that showed different trends between ER positive and ER negative groups were related to pathways involved in splicing, cell cycle, and proliferation, indicating that ER negative tumors typically evolve into highly proliferative IBTRs when compared to recurrences derived from ER positive PTs. Closer inspection of these pathways at the RNA and protein level showed enrichment of CDKs (e.g. CDK4) and the DNA replication machinery (e.g. MCM3-5; **Figure S16C-D**). The evolution of ER negative PTs into highly proliferative IBTRs might be inducing replication stress, which in turn could explain the higher mutational load in ER negative IBTRs. As the replication machinery operates, DNA repair mechanisms are responsible to correct any error or damage that might occur. Analysis of transcript/protein pairs belonging to cell cycle and DNA repair terms (source: Gene Ontology Biological Process) showed a higher expression of these genes in IBTRs derived from ER negative PTs (**Figure 6A-B**). As our previous analysis showed that *TP53* mutations was only sporadically associated to genomic, transcriptomic, or proteomic changes within our sample set (**Figure 3, S5, S8, S9, S14 and S15**), we argued that additional factors are likely involved in the accumulation of mutational features and high proliferation rates as indicated by high Ki-67 levels. The higher expression of the MYC oncogene in IBTRs derived from ER negative PTs (RNA level: fold increase 2.26, p-value < 0.001) might have been a factor in establishing high proliferation rates. However, the accumulation of genomic features (CN and SNV) only sporadically associated with absence of ER or Ki-67, suggesting that other drivers were involved in the mutational changes in these IBTRs. Consequently, we investigated the APOBEC protein family, which was previously shown to be responsible for inducing the majority of mutations in BC^49, 50^ (**Figure 6C-E** and **Figure S17**). Here we noted that APOBEC3B strongly correlated with Ki-67 levels (PT: Spearman Rho = 0.400, p-value = 0.072; IBTR: Spearman Rho = 0.674, p-value = 0.001; **Figure 6C**) and was highly expressed in ER negative PTs and IBTRs (PT: Log2Ratio = 1.728, p-value = 0.007; IBTR: Log2Ratio = 2.456, p-value < 0.001; **Figure 6D**). As APOBEC proteins are Cytosine deaminases^49^, we expected an enrichment of C>X changes in ER negative tumors, which was confirmed by a borderline enrichment of C>T transitions in this group (p-value = 0.076; **Figure 6E**). These results suggest that several factors work in parallel to enact the mutational and expression level drift of recurrent BCs from their PTs, namely the enhanced replication capacity of ER negative tumors likely driven by mechanisms outside of the ER transcriptional program (e.g. MYC) as well as the expression of mutation-inducing APOBEC proteins.

**Figure 6.**
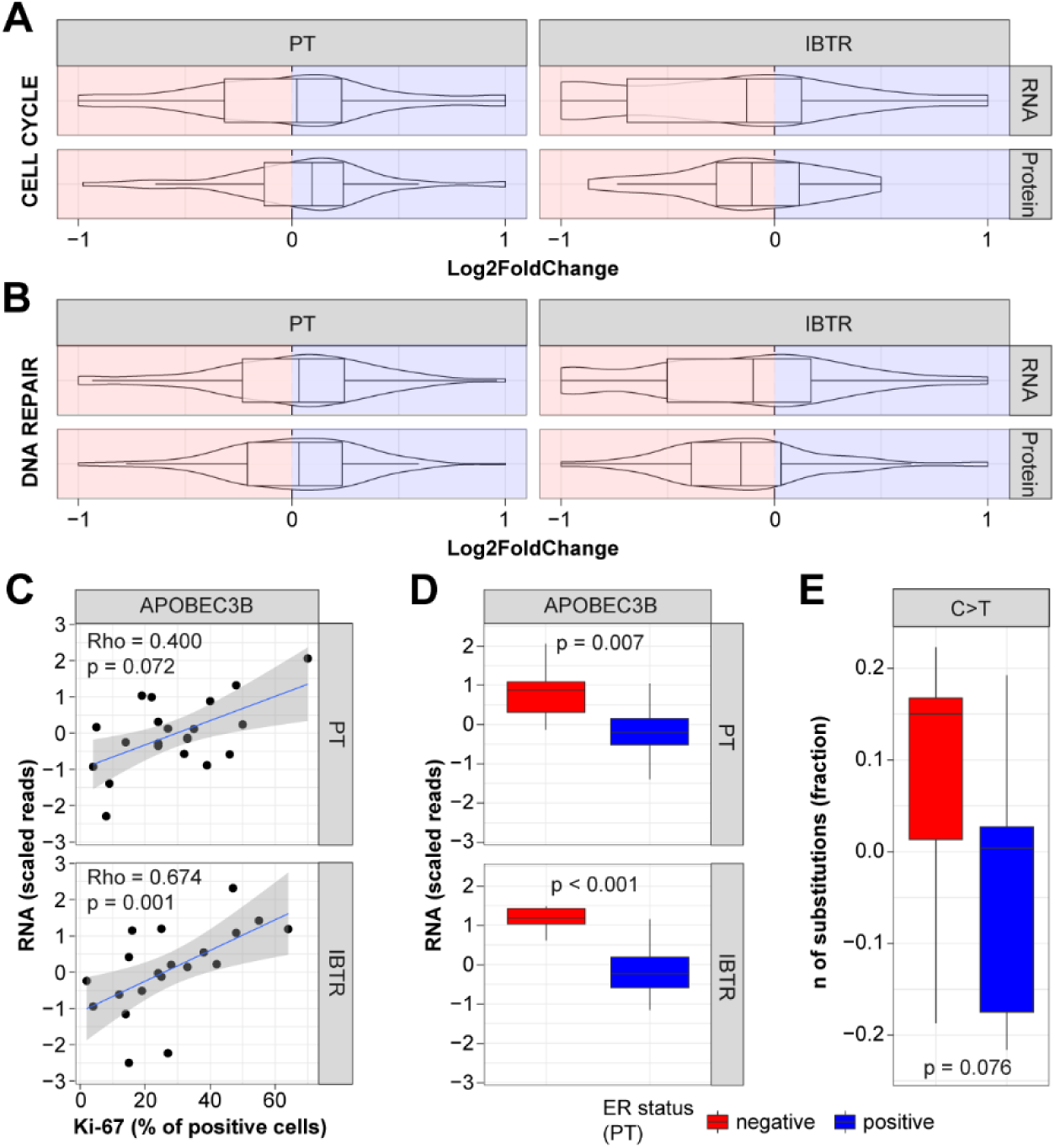
Proteogenomic factors impacting differential evolution of tumors. Multi-level analysis was performed to assess the contribution of DNA repair and APOBEC proteins to the different mutational evolution of ER positive and negative IBTRs. Panels A-B: Boxplots depicting enrichment of genes belonging to Cell Cycle and DNA repair pathways between ER positive (blue) and ER negative (red) tumors. Panel C: Correlation between APOBEC3B levels and proliferation marker Ki-67. Panel D: Differential expression of APOBEC3B genes between ER positive and ER negative tumors. Panel E: Assessment of nucleotide C-to-T transition frequency changes between ER positive and negative tumors. Acronyms: ER, estrogen receptor; Ki-67, antigen Ki-67; IBTR, ipsilateral breast tumor recurrence; PT, primary tumor.

## Discussion

IBTR is associated with an increased risk of distant metastases and breast cancer death. Molecular profiling has enabled better characterization of the mutational processes operating in BC as well as defining new therapeutic strategies and classifications schemes^12, 18, 51^. So far, most of these studies have focused on PT or DM exclusively, with limited consideration for locally recurrent BCs, which are still curable and an opportunity to define biomarkers and drug targets to prevent DM.

In this study we employed a combination of WGS, RNAseq, and MS-based proteomic analyses to elucidate the evolution of IBTRs and to define key molecular changes between the recurrence and its original matched PT. Quantitative RNA and protein analyses matched well to clinically used biomarkers, although status switches (e.g. ER) were observed (**Figure 1D**). While gain/loss of key markers is likely dependent on sub-clonal selection within the primary tumor^52, 53^, the sequencing capacity was too low to effectively reconstruct the composition and the selection of tumor sub-clones in each sample. Analysis of previously published mutational signatures^18, 19^ fitted onto our WGS data showed that the strongest contribution were from C>G and T>C enriched signatures in our samples, where signature 3 significantly associated to lack of ER expression and high proliferation in primary tumors. These results are consistent with previous observations in BC DMs^20^. In addition to this, signature 3 displayed the highest increase in IBTRs among all COSMIC signatures and was associated with hormonal receptor negative tumors, high proliferation, and radiotherapy treatment. ER negative BCs in general and triple negative BCs in particular are typically characterized by a higher degree of genomic instability than their ER positive counterparts^54, 55^. Our findings indicated that mutagenesis dependent on homologous recombination deficiency was exacerbated by radiotherapy-induced DNA damage and promoted further by high proliferation rates and subsequent activation of cell cycle checkpoints in ER negative tumors (**Figure 2D**). This relationship was confirmed in our CN and mutation evolution analyses, where a higher number of CN and mutational gains was detected in the PT-IBTR ER negative pairs (**Figure 3**). Although *TP53* mutations were enriched in the ER negative subset as previously reported^12^, we did not observe any significant association with CN or mutational gain/loss, suggesting that other mechanisms might be driving the genomic evolution of this tumor subgroup.

To assess whether the changes in genomic features impacted expression levels in a similar fashion, paired PT-IBTR distances were calculated based on CN, mutation, RNA expression, and protein abundance (**Figure 4**). Interestingly, we here found that changes at the CN and mutational level often impacted transcript and protein abundance, with positive correlations between genomic, transcriptomic, and proteomic distances. On top of this, our analyses revealed that lack of hormonal receptors and proliferation rates implied a drift not only within the space of genomic features, but also a reprogramming of both the transcriptome and the proteome. Given the fact that ER positive and negative PTs display different transcriptional programs^56, 57^ and often feature different sets of driver mutations (e.g. *PIK3CA vs TP53*)^12^, it is reasonable to believe these features have an effect on cancer evolution. Differential gene/protein expression and pathway analyses within the ER positive and ER negative recurrence-forming PTs revealed two differential evolutionary routes, where ER negative IBTRs were enriched in cell cycle, DNA replication, and transcription, while ER positive tumors were geared toward metabolic pathways (ER positive; **Figure 5**). In addition to cell cycle-related genes, an enrichment of APOBEC3B was also detected in ER negative tumors (**Figure 6**). APOBEC3B is a known cancer mutagen often overexpressed in BC and seemingly responsible for ∼80% of the mutational load in these tumors^49, 50^. APOBEC3B action in breast cancer has been shown to change in relation to the expression of ER, of which is an interactor recruited at binding sites, promoting DNA strand breaks^58^. This synergistic action is responsible for poor clinical outcomes in ER positive BCs^58–60^. While ER negative tumors have been reported to express high levels of APOBEC3B^59^, this has not been linked to clinical outcome nor have its effect on the mutational landscape of these tumor subset been characterized. ER negative tumors are generally indicative of poor prognosis due to the fact that multiple mechanisms are enacted to enable tumor cell proliferation outside of the ER transcriptional program^12, 14, 57^, conferring new features to cancer cells such tissue invasion^61^ or immune evasion^62^. In addition, several studies have shown that ER negative and triple-negative BC constitute a molecularly heterogeneous group^63–66^. In the light of this, the role and clinical association of APOBEC family members might be either concomitant to other factors, hence the non-significant contribution of APOBEC-related signatures (i.e. signature 2 and 13) in this subset, or confounded by the numerous processes at work in these tumors.

Investigation of the role of APOBEC family members in ER negative BCs would entice the analysis of subtype-stratified cohorts to better define their relationship with clinical outcomes, mutational processes, and other key factors (e.g. immune system). Further mechanistic studies assessing the interaction of APOBEC enzymes with cancer drivers (e.g. MYC) or other factors active in ER negative cancers would allow to quantify the impact on these tumorś mutational landscape and define new drug targets or alternative treatment regimens, such as in the case of PARP1-inhibitor mediated synthetic lethality^67^.

While our study could not recapitulate IBTR features due to low power or resolve tumor clonal evolution at high resolution due to shallow sequencing, it shows how the mutational landscape of recurrent breast cancers diversifies based on the expression of hormonal receptors, with repercussions at the transcriptome and proteome levels and repurposing the cell machinery towards DNA replication and proliferation, indicating these mechanisms should be targeted to prevent IBTR formation.

## Data availability

The sequencing data (WGS and RNAseq) presented in this study are available upon request from the corresponding authors, and after additional ethical approval. The data are not publicly available due to ethical considerations.

DIA MS data, and their respective search result files have been deposited to the ProteomeXchange Consortium via the PRIDE partner repository^68^ with the dataset identifier: PXD032266.

## Acknowledgements

We gratefully thank Sara Baker, Carina Forsare, Kristina Lövgren and Anna-Lena Borg for excellent technical assistance. We also thank the biobanks of the South Sweden Breast Cancer Group (SSBCG), the Biobank at the Department of Oncology and Pathology Lund University biobank at Cancer Center Karolinska and the Biobank at Academic Hospital in Uppsala and Department of Pathology, Uppsala University, for collecting the samples and making them available for studies. Sequencing was performed by the SNP&SEQ Technology Platform in Uppsala. The facility is part of the National Genomics Infrastructure (NGI) Sweden and Science for Life Laboratory. The SNP&SEQ Platform is also supported by the Swedish Research Council and the Knut and Alice Wallenberg Foundation. Parts of the computational analysis was performed on resources provided by SNIC through Uppsala Multidisciplinary Center for Advanced Computational Science (UPPMAX) under Project sens2016010 and the authors would like to acknowledge support from Science for Life Laboratory, the National Genomics Infrastructure (NGI), National Bioinformatics Infrastructure Sweden (NBIS) and UPPMAX for providing assistance in massive parallel sequencing and computational infrastructure.

## Author contribution

TDM, PP, MS, JM and EN designed the study. TDM, MS, PP, SR, and SL processed sequencing data. TDM and PP processed proteomic data. LT and GP performed and scored immunohistochemical stainings. MS, FW, IF, PM, LM, JM, MF, and EN administered medical records, provided analysis platforms and support. TDM wrote the manuscript with assistance from all authors.

## Abbreviations

AF: allelic frequency
BC: breast cancer
CN: copy number
DIA: data independent acquisition
DM: distant metastasis
ER: estrogen receptor
ERBB2/Her2: receptor tyrosine-protein kinase erbB-2
FA: formic acid
FFPE: formalin-fixed and paraffin-embedded
GSEA: gene set enrichment analysis
Ki-67: antigen Ki-67
IBTR: ipsilateral breast tumor recurrence
IBTRFS: ipsilateral breast tumor recurrence-free survival
IHC: immunohistochemistry
IQR: interquartile range
MS: mass spectrometry
NBIS: National bioinformatics infrastructure Sweden
NGI: national genomics infrastructure
PgR: progesterone receptor
PoN: panel of normal
PT: primary tumor
RNAseq: RNA sequencing
SNV: single nucleotide variant
SSBCG: South Sweden Breast Cancer Group
TP53: tumor protein p53
UPPMAX: Uppsala multidisciplinary center for advanced computational science
WGS: whole genome sequencing.

## Notes

### Competing Interest Statement

PM and MF research contract with PFS Genomics

