## Supplementary Figures 1-17 for "”Evolution of ipsilateral breast cancer decoded by proteogenomics”"

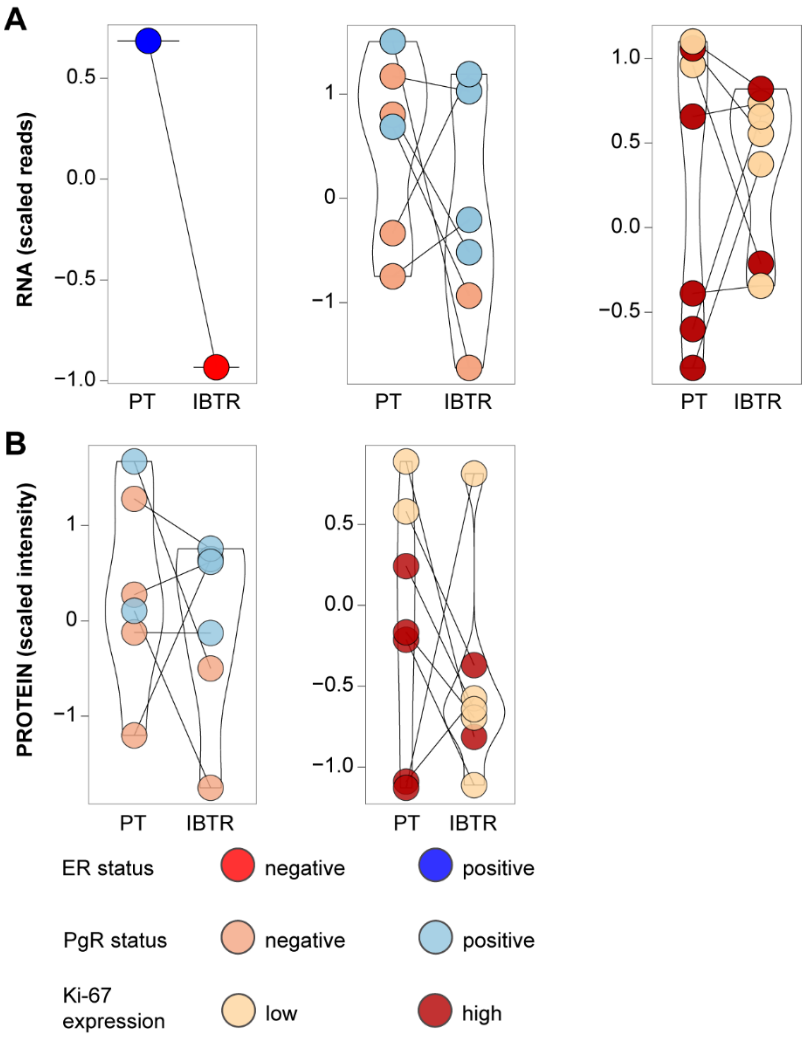


**Figure S1 – Validation of biomarker switch by RNA and proteomics.** Changes in the status of ER, PgR, and Ki-67 were detected when analyzing PT-IBTR pairs by IHC. Panels A and B depict RNA and protein measurement of the three markers, respectively.

Acronyms: ER, estrogen receptor; Ki-67, antigen Ki-67; IBTR: ipsilateral breast tumor recurrence; IHC: immunohistochemistry; PgR, progesterone receptor; PT, primary tumor.

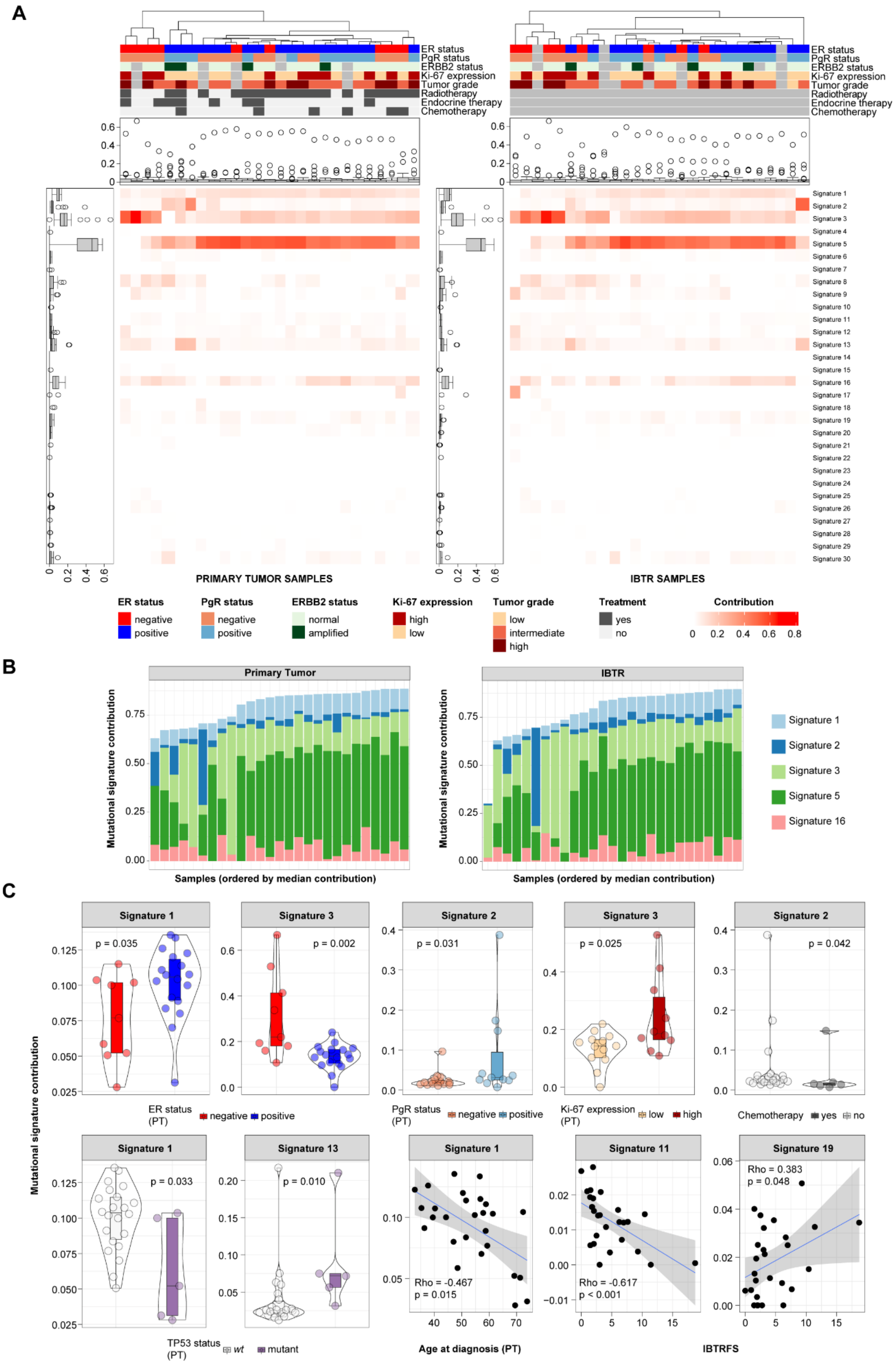
A
ER status
PgR status
ERBB2 status
Ki-67 expression
Tumor grade
Radiotherapy
Endocrine therapy
Chemotherapy
PRIMARY TUMOR SAMPLES
IBTR SAMPLES
Signature 1
Signature 2
Signature 3
Signature 4
Signature 5
Signature 6
Signature 7
Signature 8
Signature 9
Signature 10
Signature 11
Signature 12
Signature 13
Signature 14
Signature 15
Signature 16
Signature 17
Signature 18
Signature 19
Signature 20
Signature 21
Signature 22
Signature 23
Signature 24
Signature 25
Signature 26
Signature 27
Signature 28
Signature 29
Signature 30
ER status: negative, positive
PgR status: negative, positive
ERBB2 status: normal, amplified
Ki-67 expression: high, low
Tumor grade: low, intermediate, high
Treatment: yes, no
Contribution: 0, 0.2, 0.4, 0.6, 0.8
B
Primary Tumor
IBTR
Mutational signature contribution
Samples (ordered by median contribution)
Signature 1
Signature 2
Signature 3
Signature 5
Signature 16
C
Mutational signature contribution
Signature 1, p = 0.035
Signature 3, p = 0.002
ER status (PT): negative, positive
Signature 2, p = 0.031
PgR status (PT): negative, positive
Signature 3, p = 0.025
Ki-67 expression (PT): low, high
Signature 2, p = 0.042
Chemotherapy: yes, no
Signature 1, p = 0.033
Signature 13, p = 0.010
TP53 status (PT): wt, mutant
Signature 1, Rho = -0.467, p = 0.015
Age at diagnosis (PT)
Signature 11, Rho = -0.617, p < 0.001
Signature 19, Rho = 0.383, p = 0.048
IBTRFS

**Figure S2 - Mutational signature contribution within the sample cohort.** COSMIC-derived mutational signatures were fitted into our dataset and analyzed for their overall contribution in all samples. Panel A illustrates side-by-side heatmaps of mutational signature contribution across PT and IBTR samples. Boxplot to the side and top of each heatmap recapitulate signature contribution over the dataset and in each sample, respectively. Panel B depicts bar charts of top 5 contributing signatures (selected based on median contribution over the dataset). Panel C shows significant results out of the association of clinical variables to mutational signature contribution within the primary tumor subset.

Acronyms: ER, estrogen receptor; Ki-67, antigen Ki-67; IBTR: ipsilateral breast tumor recurrence; IBTRFS, IBTR-free survival; PgR, progesterone receptor; PT, primary tumor; TP53, tumor protein p53.

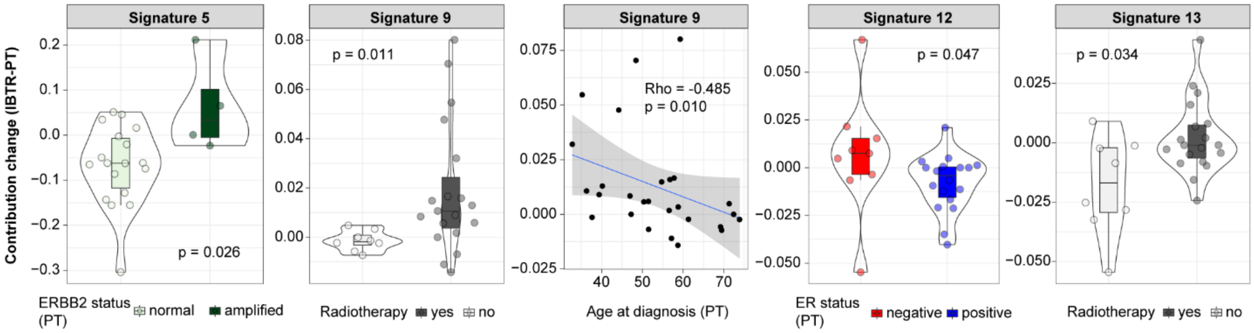


**Figure S3 - Evolution of mutational processes.** We evaluated the contribution of the 30 COSMIC mutational signatures within our samples (PT and IBTR subsets). Contribution delta was then calculated as a measure of mutational process evolution for each PT-IBTR pair. Significant associations between changes in mutational signature contribution and clinical variables is shown.

Acronyms: ER, estrogen receptor; ERBB2/Her2, receptor tyrosine-protein kinase erbB-2; IBTR: ipsilateral breast tumor recurrence; PT, primary tumor.

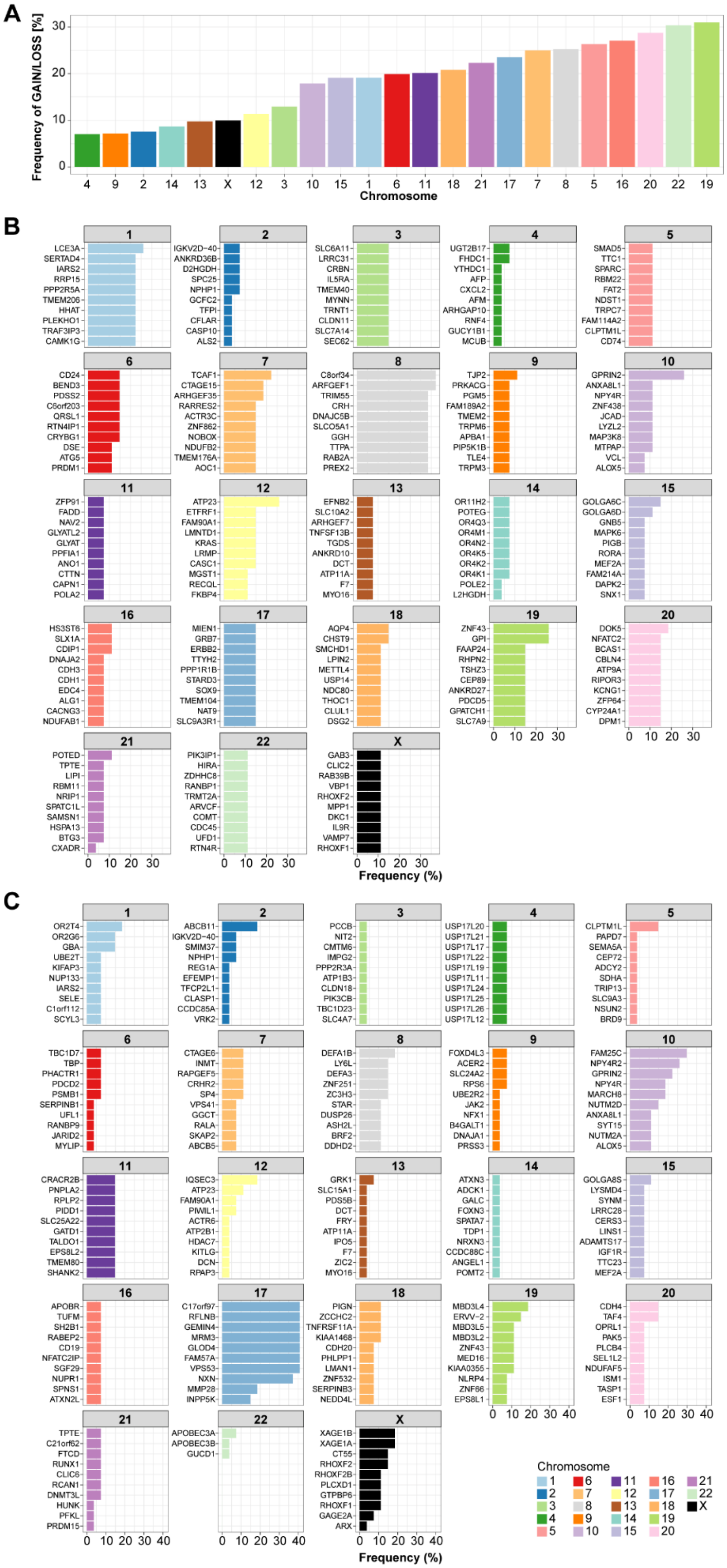

A
Frequency of GAIN/LOSS [%]
0 10 20 30
4 9 2 14 13 X 12 3 10 15 1 6 11 18 21 17 7 8 5 16 20 22 19
Chromosome
B
1: LCE3A SERTAD4 IARS2 RRP15 PPP2R5A TMEM206 HHAT PLEKHO1 TRAF3IP3 CAMK1G
2: IGKV2D−40 ANKRD36B D2HGDH SPC25 NPHP1 GCFC2 TFPI CFLAR CASP10 ALS2
3: SLC6A11 LRRC31 CRBN IL5RA TMEM40 MYNN TRNT1 CLDN11 SLC7A14 SEC62
4: UGT2B17 FHDC1 YTHDC1 AFP CXCL2 AFM ARHGAP10 RNF4 GUCY1B1 MCUB
5: SMAD5 TTC1 SPARC RBM22 FAT2 NDST1 TRPC7 FAM114A2 CLPTM1L CD74
6: CD24 BEND3 PDSS2 C6orf203 QRSL1 RTN4IP1 CRYBG1 DSE ATG5 PRDM1
7: TCAF1 CTAGE15 ARHGEF35 RARRES2 ACTR3C ZNF862 NOBOX NDUFB2 TMEM176A AOC1
8: C8orf34 ARFGEF1 TRIM55 CRH DNAJC5B SLCO5A1 GGH TTPA RAB2A PREX2
9: TJP2 PRKACG PGM5 FAM189A2 TMEM2 TRPM6 APBA1 PIP5K1B TLE4 TRPM3
10: GPRIN2 ANXA8L1 NPY4R ZNF438 JCAD LYZL2 MAP3K8 MTPAP VCL ALOX5
11: ZFP91 FADD NAV2 GLYATL2 GLYAT PPFIA1 ANO1 CTTN CAPN1 POLA2
12: ATP23 ETFRF1 FAM90A1 LMNTD1 KRAS LRMP CASC1 MGST1 RECQL FKBP4
13: EFNB2 SLC10A2 ARHGEF7 TNFSF13B TGDS ANKRD10 DCT ATP11A F7 MYO16
14: OR11H2 POTEG OR4Q3 OR4M1 OR4N2 OR4K5 OR4K2 OR4K1 POLE2 L2HGDH
15: GOLGA6C GOLGA6D GNB5 MAPK6 PIGB RORA MEF2A FAM214A DAPK2 SNX1
16: HS3ST6 SLX1A CDIP1 DNAJA2 CDH3 CDH1 EDC4 ALG1 CACNG3 NDUFAB1
17: MIEN1 GRB7 ERBB2 TTYH2 PPP1R1B STARD3 SOX9 TMEM104 NAT9 SLC9A3R1
18: AQP4 CHST9 SMCHD1 LPIN2 METTL4 USP14 NDC80 THOC1 CLUL1 DSG2
19: ZNF43 GPI FAAP24 RHPN2 TSHZ3 CEP89 ANKRD27 PDCD5 GPATCH1 SLC7A9
20: DOK5 NFATC2 BCAS1 CBLN4 ATP9A RIPOR3 KCNG1 ZFP64 CYP24A1 DPM1
21: POTED TPTE LIPI RBM11 NRIP1 SPATC1L SAMSN1 HSPA13 BTG3 CXADR
22: PIK3IP1 HIRA ZDHHC8 RANBP1 TRMT2A ARVCF COMT CDC45 UFD1 RTN4R
X: GAB3 CLIC2 RAB39B VBP1 RHOXF2 MPP1 DKC1 IL9R VAMP7 RHOXF1
0 10 20 30
Frequency (%)
C
1: OR2T4 OR2G6 GBA UBE2T KIFAP3 NUP133 IARS2 SELE C1orf112 SCYL3
2: ABCB11 IGKV2D−40 SMIM37 NPHP1 REG1A EFEMP1 TFCP2L1 CLASP1 CCDC85A VRK2
3: PCCB NIT2 CMTM6 IMPG2 PPP2R3A ATP1B3 CLDN18 PIK3CB TBC1D23 SLC4A7
4: USP17L20 USP17L21 USP17L17 USP17L22 USP17L19 USP17L11 USP17L24 USP17L25 USP17L26 USP17L12
5: CLPTM1L PAPD7 SEMA5A CEP72 ADCY2 SDHA TRIP13 SLC9A3 NSUN2 BRD9
6: TBC1D7 TBP PHACTR1 PDCD2 PSMB1 SERPINB1 UFL1 RANBP9 JARID2 MYLIP
7: CTAGE6 INMT RAPGEF5 CRHR2 SP4 VPS41 GGCT RALA SKAP2 ABCB5
8: DEFA1B LY6L DEFA3 ZNF251 ZC3H3 STAR DUSP26 ASH2L BRF2 DDHD2
9: FOXD4L3 ACER2 SLC24A2 RPS6 UBE2R2 JAK2 NFX1 B4GALT1 DNAJA1 PRSS3
10: FAM25C NPY4R2 GPRIN2 NPY4R MARCH8 NUTM2D ANXA8L1 SYT15 NUTM2A ALOX5
11: CRACR2B PNPLA2 RPLP2 PIDD1 SLC25A22 GATD1 TALDO1 EPS8L2 TMEM80 SHANK2
12: IQSEC3 ATP23 FAM90A1 PIWIL1 ACTR6 ATP2B1 HDAC7 KITLG DCN RPAP3
13: GRK1 SLC15A1 PDS5B DCT FRY ATP11A IPO5 F7 ZIC2 MYO16
14: ATXN3 ADCK1 GALC FOXN3 SPATA7 TDP1 NRXN3 CCDC88C ANGEL1 POMT2
15: GOLGA8S LYSMD4 SYNM LRRC28 CERS3 LINS1 ADAMTS17 IGF1R TTC23 MEF2A
16: APOBR TUFM SH2B1 RABEP2 CD19 NFATC2IP SGF29 NUPR1 SPNS1 ATXN2L
17: C17orf97 RFLNB GEMIN4 MRM3 GLOD4 FAM57A VPS53 NXN MMP28 INPP5K
18: PIGN ZCCHC2 TNFRSF11A KIAA1468 CDH20 PHLPP1 LMAN1 ZNF532 SERPINB3 NEDD4L
19: MBD3L4 ERVV−2 MBD3L5 MBD3L2 ZNF43 MED16 KIAA0355 NLRP4 ZNF66 EPS8L1
20: CDH4 TAF4 OPRL1 PAK5 PLCB4 SEL1L2 NDUFAF5 ISM1 TASP1 ESF1
21: TPTE C21orf62 FTCD RUNX1 CLIC6 RCAN1 DNMT3L HUNK PFKL PRDM15
22: APOBEC3A APOBEC3B GUCD1
X: XAGE1B XAGE1A CT55 RHOXF2 RHOXF2B PLCXD1 GTPBP6 RHOXF1 GAGE2A ARX
0 10 20 30 40
Frequency (%)
Chromosome
1 2 3 4 5 6 7 8 9 10 11 12 13 14 15 16 17 18 19 20 21 22 X

**Figure S4 - Frequency of copy number changes between primary and recurrent tumors.** Overview of CN changes (i.e. GAIN/LOSS) between matched samples stratified by chromosome. Panel A: Frequency of CN changes between PT and IBTR samples. Panel B-C: Top 10 gene-per-chromosome gains (B) and losses (C) in IBTR tumors.

Acronyms: CN, copy number; IBTR: ipsilateral breast tumor recurrence; PT, primary tumor.

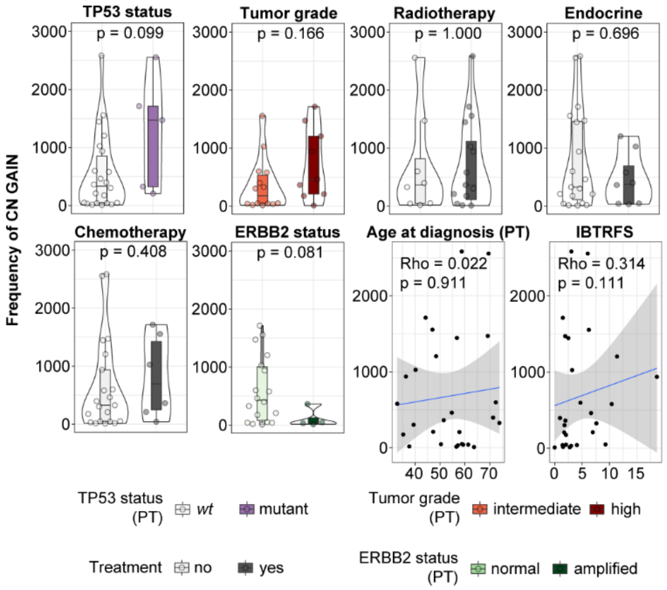


**Figure S5 – Relationship between copy number gains and clinical variables.** Panels depict association between CN gain in IBTR samples to clinical and histopathological features, and therapeutic intervention.

Acronyms: CN, copy number; ERBB2/Her2, receptor tyrosine-protein kinase erbB-2; IBTR, ipsilateral breast tumor recurrence; IBTRFS, IBTR-free survival; PT, primary tumor; TP53, tumor protein p53.

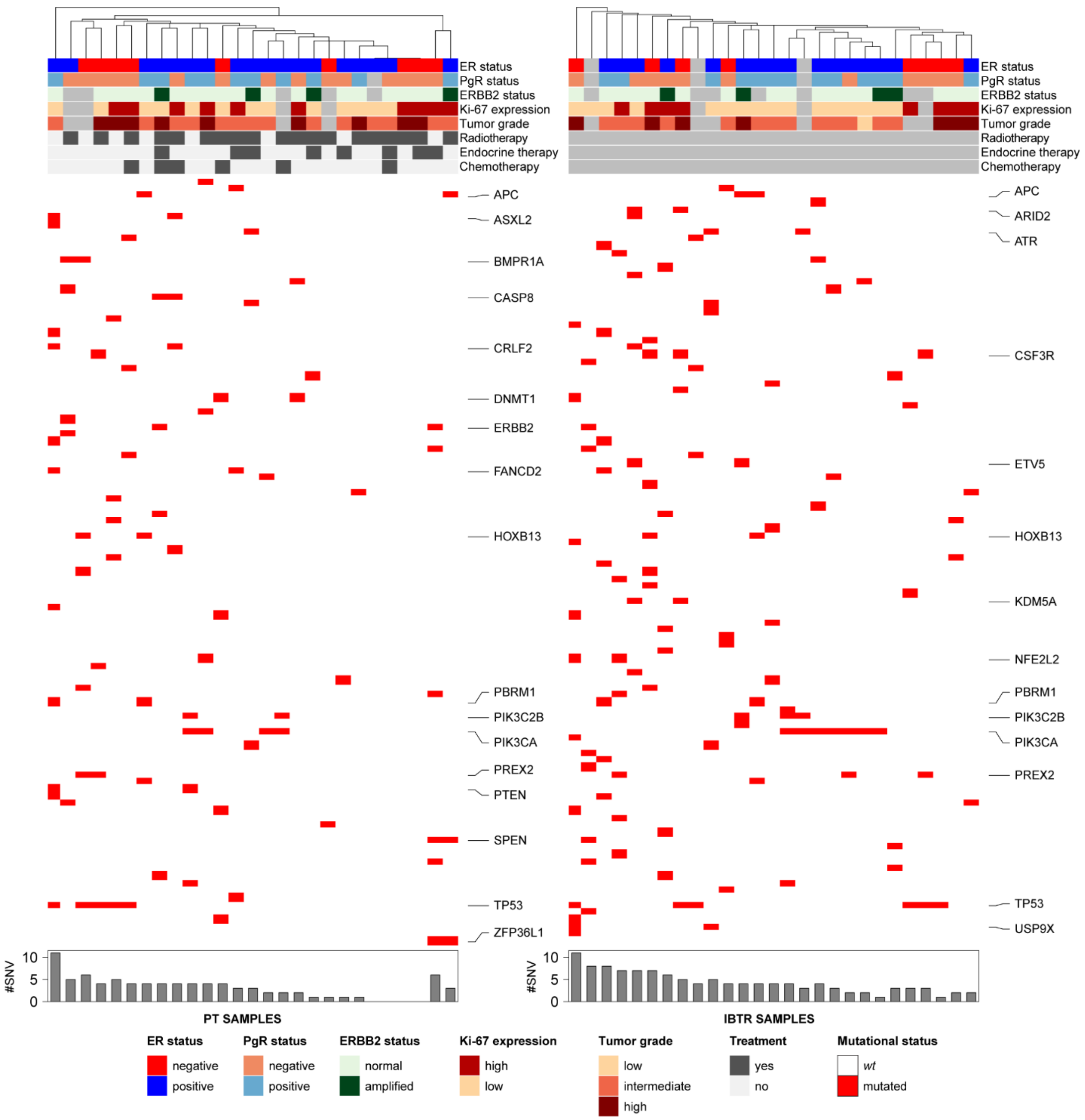


**Figure S6 – Changes in copy number and key drivers.** SNV status of COSMIC cancer genes was evaluated in our cohort and reported for each subset (PT and IBTR) as heatmaps (light gray boxes represent missing values). Bottom bar charts display sample-wise frequencies of SNVs.

Acronyms: ER, estrogen receptor; ERBB2/Her2, receptor tyrosine-protein kinase erbB-2; Ki-67, antigen Ki-67; IBTR, ipsilateral breast tumor recurrence; PgR, progesterone receptor; PT, primary tumor; SNV, single nucleotide variant.

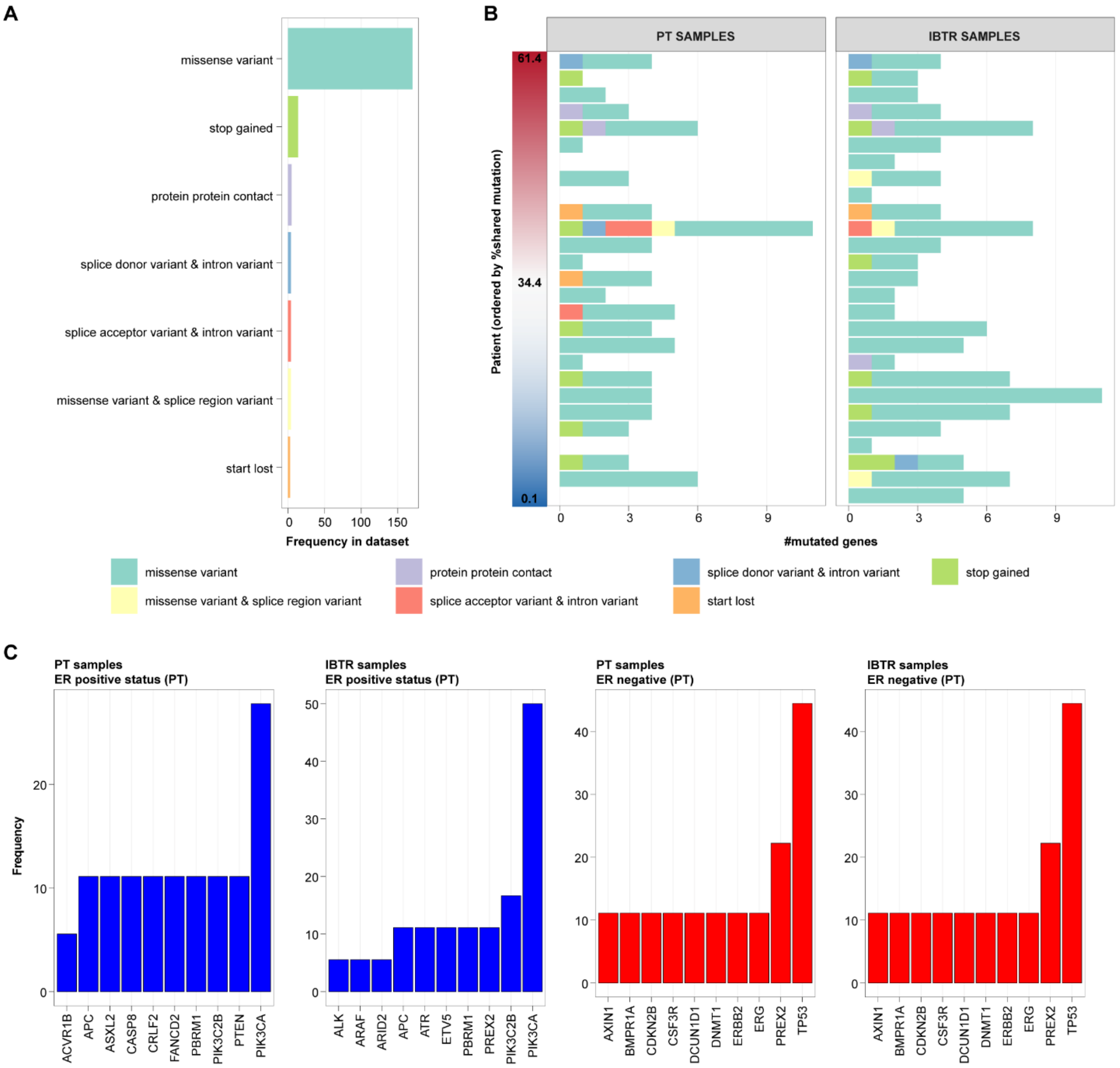


**Figure S7 - Overview of mutational features.** Filtered SNV from key cancer genes were analyzed to assess their frequency and their changes across the dataset and between matched pairs. Panel A: Mutational frequency in the dataset by type. Panel B: Mutational frequency in each paired sample. Panel C: Top 10 mutated genes in by ER status in primary and recurrent tumors.

Acronyms: ER, estrogen receptor; IBTR, ipsilateral breast tumor recurrence; PT, primary tumor; SNV, single nucleotide variant.

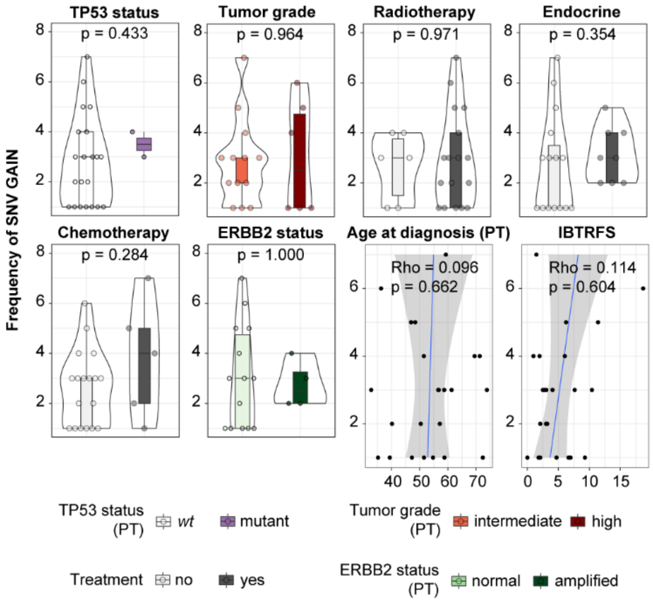


**Figure S8 – Relationship SNV gains to clinical variables.** Panels depict association between SNV gain in IBTR samples to clinical and histopathological features, and therapeutic intervention.

Acronyms: CN, copy number; ERBB2/Her2, receptor tyrosine-protein kinase erbB-2; IBTR, ipsilateral breast tumor recurrence; IBTRFS, IBTR-free survival; PT, primary tumor; TP53, tumor protein p53.

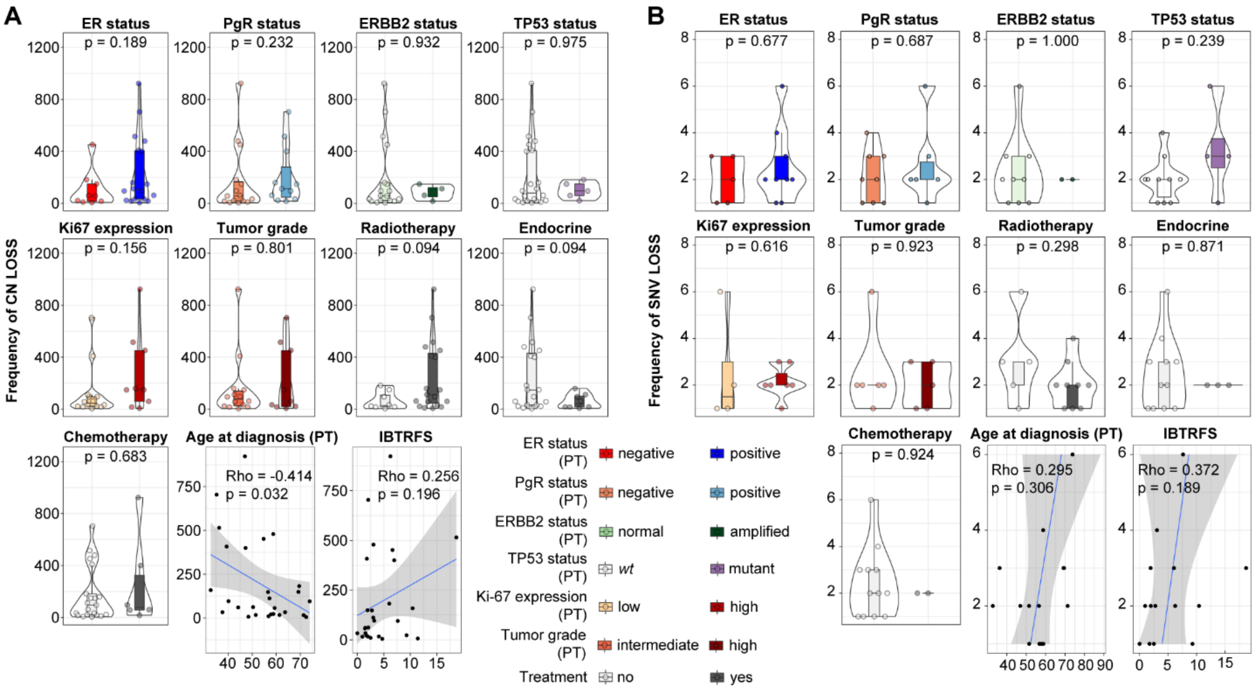


**Figure S9 – Relationship between copy number and mutational losses to clinical variables.** Panels A-B: Association of frequencies of CN (A) and SNV (B) loss between PTs and IBTRs to clinical and histopathological features, and therapeutic intervention.

Acronyms: CN, copy number; ER, estrogen receptor; ERBB2/Her2, receptor tyrosine-protein kinase erbB-2; Ki-67, antigen Ki-67; IBTR, ipsilateral breast tumor recurrence; IBTRFS, IBTR-free survival; PgR, progesterone receptor; PT, primary tumor; SNV, single nucleotide variant; TP53, tumor protein p53.

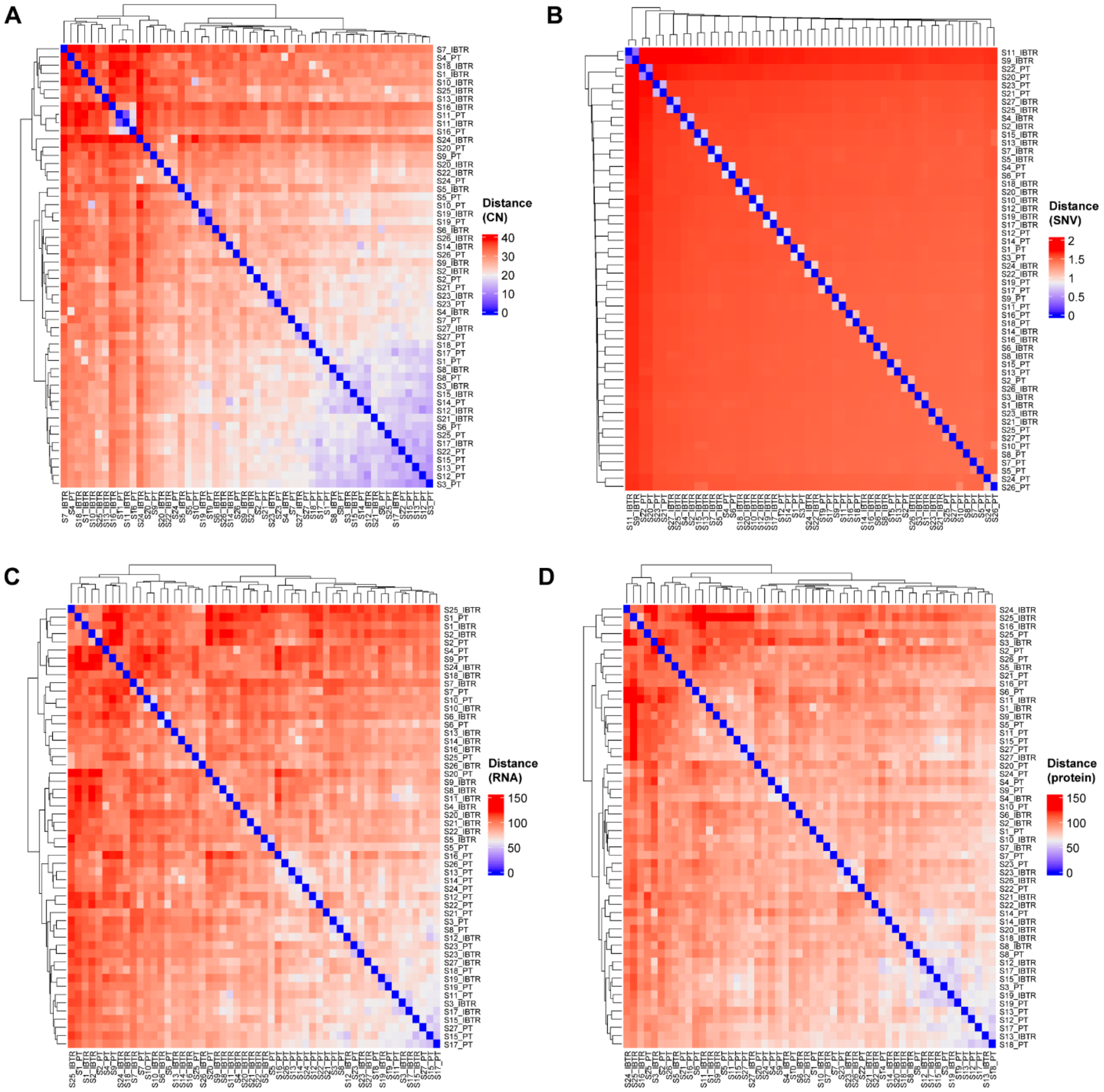


**Figure S10 – Distance matrices between samples across omics levels.** Panels display clustered heatmaps of Euclidean distance matrices computed between all samples (PTs and IBTRs) at the CN (A), SNV (B), RNA (C), and protein (D) levels. Distances were computed for matching gene-transcript-protein at the CN, RNA, and protein levels.

Acronyms: CN, copy number; IBTR, ipsilateral breast tumor recurrence; PT, primary tumor; SNV, single nucleotide variant.

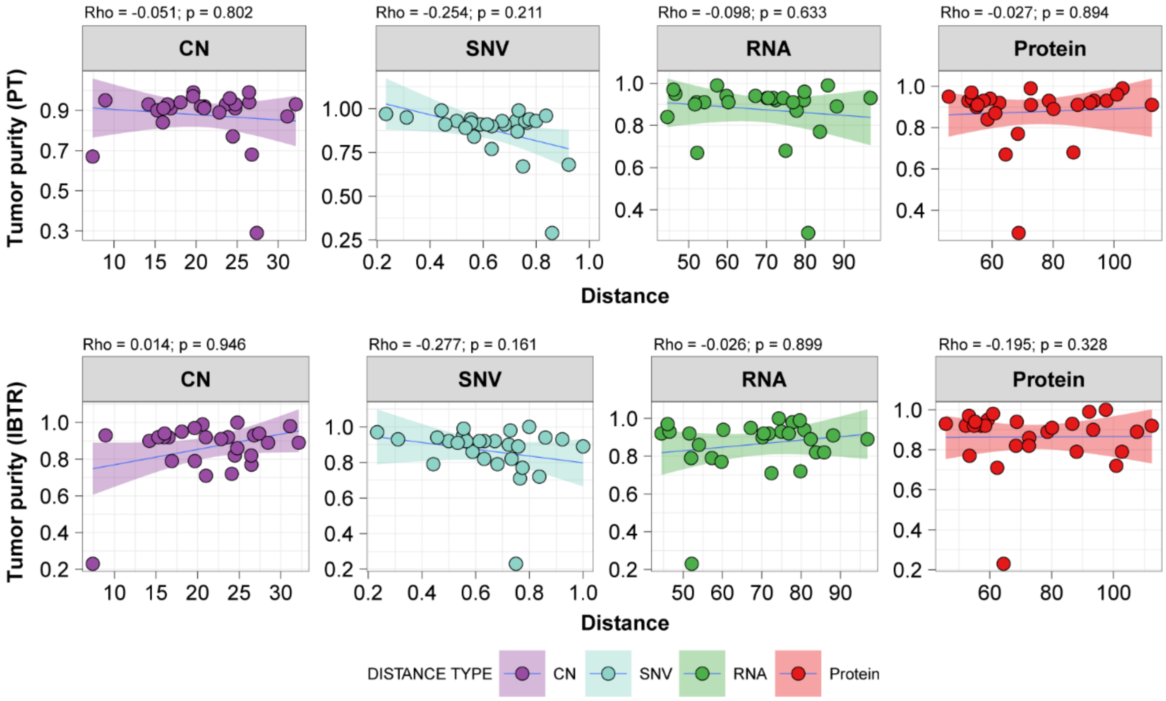


**Figure S11 – Relationship between distances and tumor purity.** Association between distance and TPES-based calculation of tumor purity (see Methods) was plotted against distances of all omics layers.

Acronyms: CN, copy number; IBTR, ipsilateral breast tumor recurrence; PT, primary tumor; SNV, single nucleotide variant.

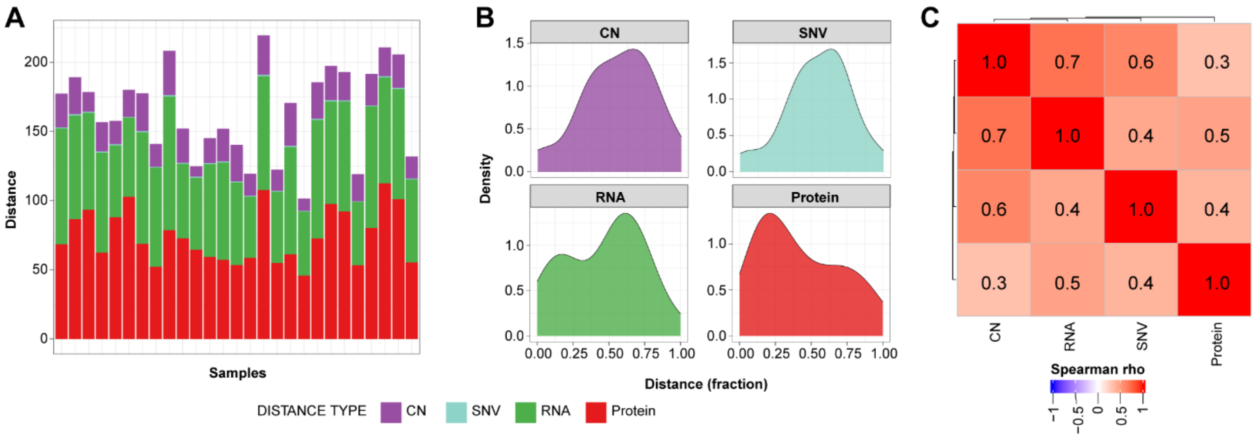


**Figure S12 – Distances between PT-IBTR pairs.** Euclidean distances were calculated between tumor pairs at the CN, SNV, transcript, and protein levels as a measure of evolutionary drift. Panel A: stacked bar chart of distances for each sample. Panel B: density plots of PT-IBTR distances (expressed as fractions) for each data layer. Panel C: Spearman correlation matrix of distances (CN, mutational, RNA, and protein).

Acronyms: CN, copy number; IBTR, ipsilateral breast tumor recurrence; PT, primary tumor; SNV, single nucleotide variant.

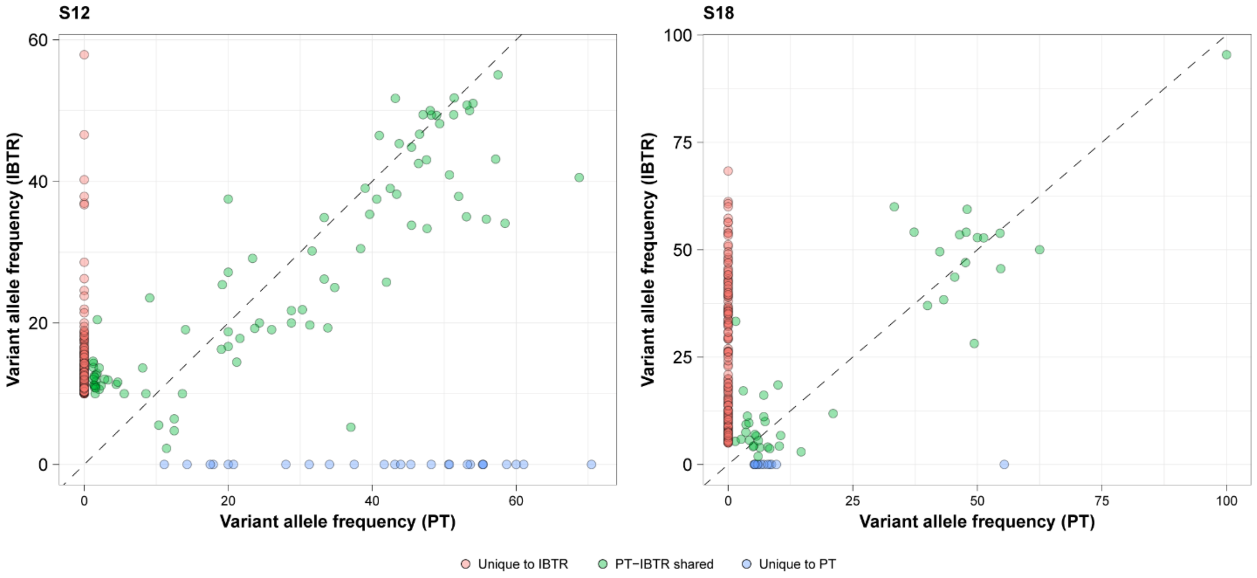


**Figure S13 – Comparison of PT-IBTR clonal composition.** Scatter plots show variant allele frequencies of identified variants in PT and IBTR pairs.

Acronyms: IBTR, ipsilateral breast tumor recurrence; PT, primary tumor.

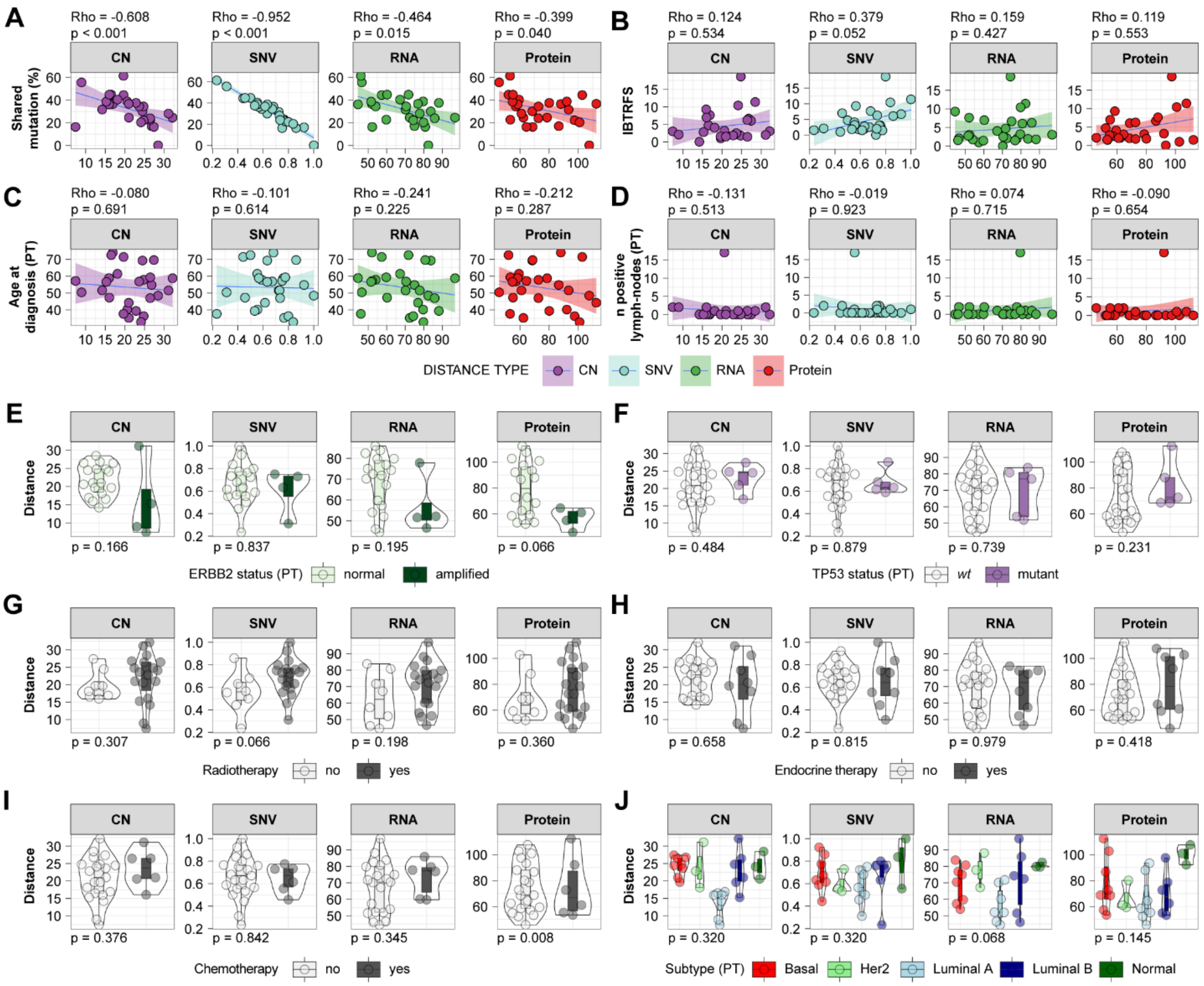


**Figure S14 – Relationship between distances and primary tumor clinical variables.** Association between distance and clinical variables of PT samples.

Acronyms: CN, copy number; ER, estrogen receptor; ERBB2/Her2, receptor tyrosine-protein kinase erbB-2; Ki-67, antigen Ki-67; IBTR, ipsilateral breast tumor recurrence; IBTRFS, IBTR-free survival; PgR, progesterone receptor; SNV, single nucleotide variant; TP53, tumor protein p53.

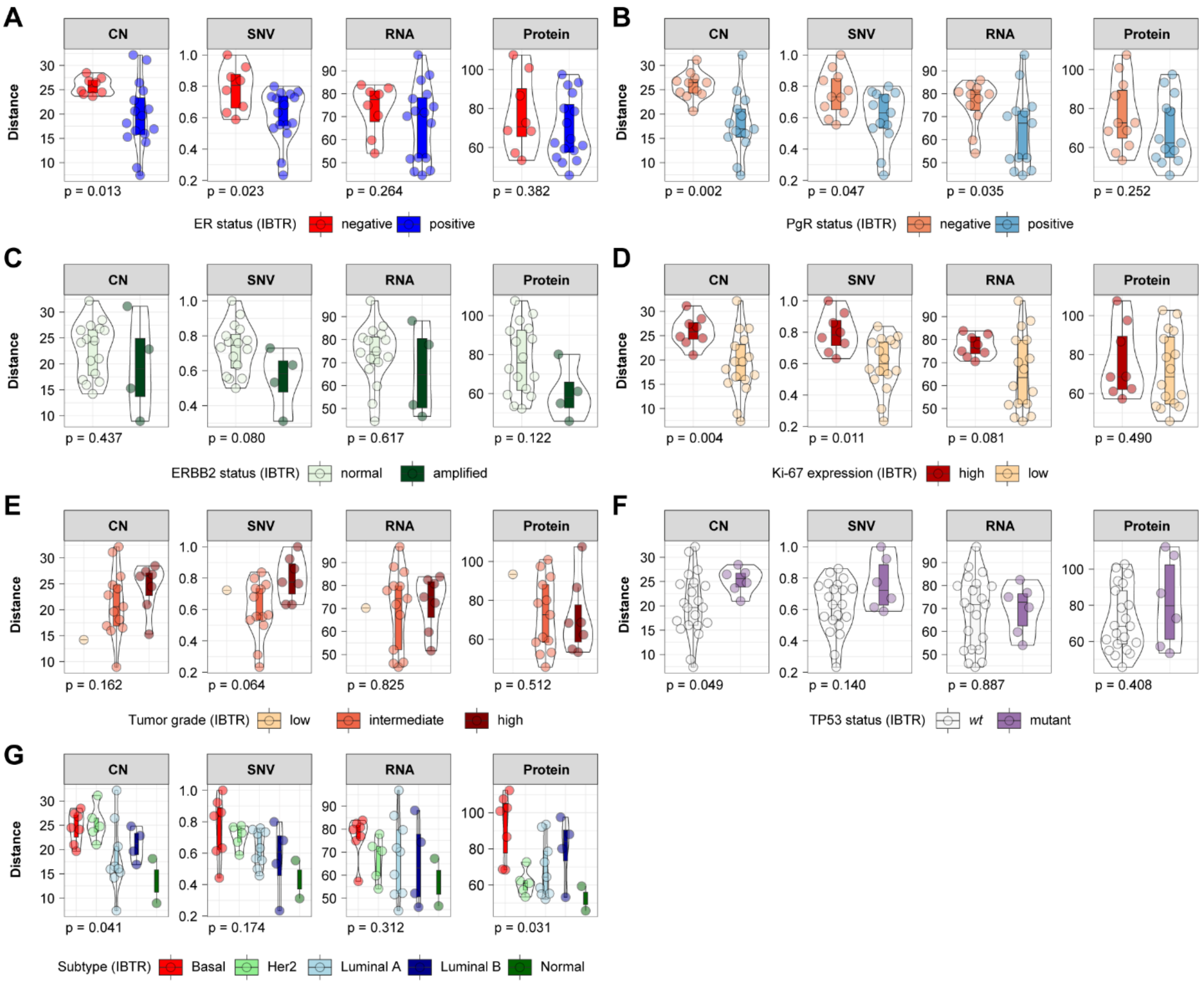


**Figure S15 – Relationship between distances and recurrent tumor clinical variables.** Association between distance and clinical variables of IBTR samples.

Acronyms: CN, copy number; ER, estrogen receptor; ERBB2/Her2, receptor tyrosine-protein kinase erbB-2; Ki-67, antigen Ki-67; IBTR, ipsilateral breast tumor recurrence; PgR, progesterone receptor; SNV, single nucleotide variant; TP53, tumor protein p53.

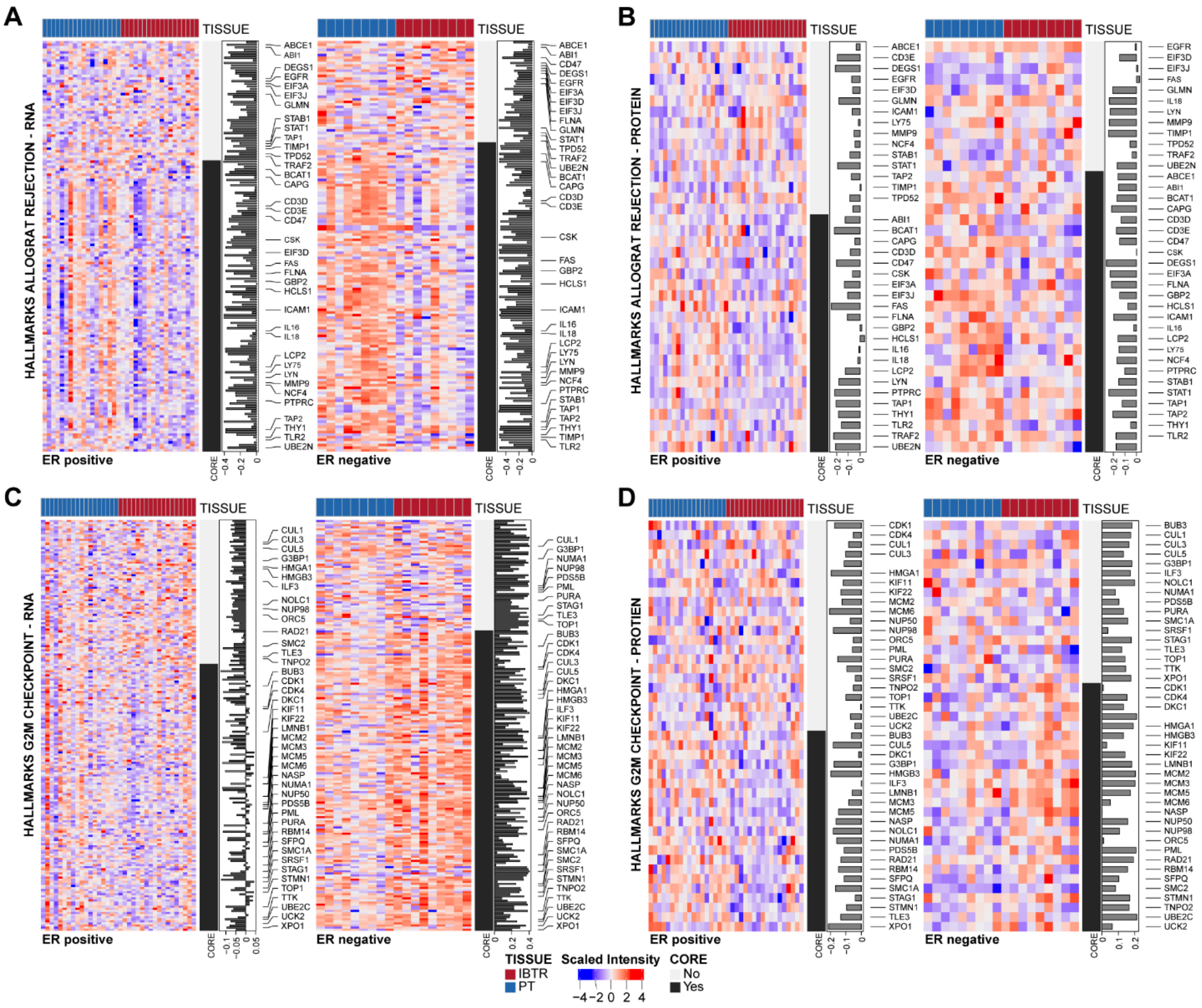


**Figure S16 – Key differentially enriched pathways between ER positive and ER negative PT-IBTR groups.** Panels A-B: Pathway enrichment divergence in ER positive and negative patients at the RNA and protein levels. Panels C-D: Differences at the transcript and protein levels within immune signaling and proliferation pathways.

Acronyms: ER, estrogen receptor; IBTR, ipsilateral breast tumor recurrence; PT, primary tumor.

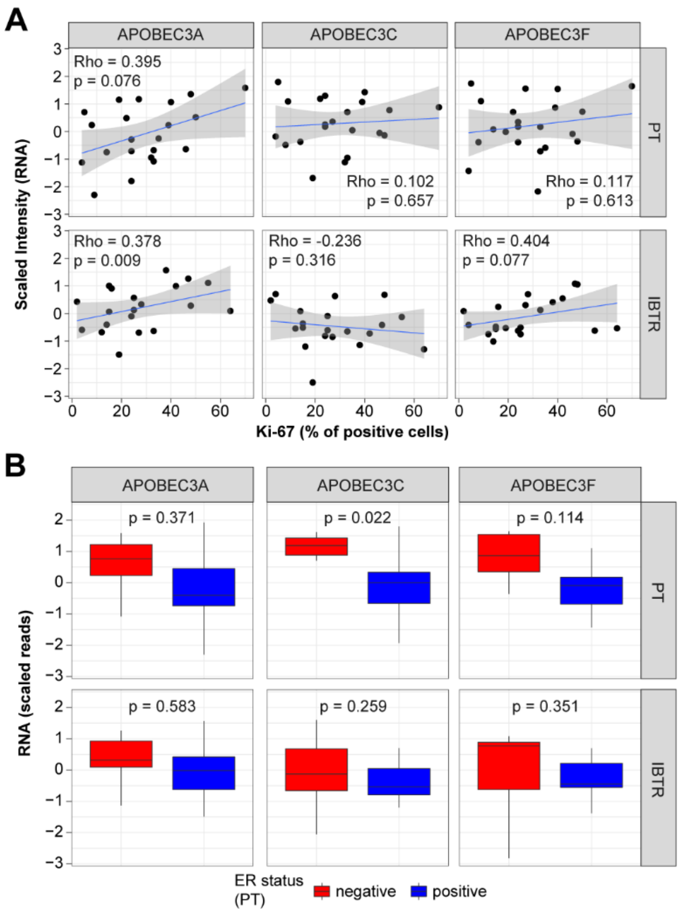


**Figure S17 – Relationship between the APOBEC family, ER status, and Ki-67 levels.** Panel A: Correlation between APOBEC levels and proliferation marker Ki-67. Panel B: Differential expression of APOBEC genes between ER positive and ER negative tumors.

Acronyms: ER, estrogen receptor; Ki-67, antigen Ki-67; IBTR, ipsilateral breast tumor recurrence; PT, primary tumor.
